# Postharvest tomato shelf life is genetically distinct from fruit firmness: evidence from two F_2_ populations

**DOI:** 10.64898/2026.07.22.739999

**Authors:** Carlo Rosati, Fabien Tirado, Giuseppe Aprea, Frédérique Bitton, Marie Brault, Renaud Duboscq, Paola Ferrante, Karine Pellegrino, Catia Stamigna, Giovanni Giuliano, Mathilde Causse

## Abstract

Reducing postharvest fruit loss without compromising fruit quality is a major goal in tomato breeding. Fruit shelf life is a complex trait, influenced by postharvest changes in fruit firmness and weight, as well as by both genetic and environmental factors. A QTL mapping experiment was conducted to identify loci associated with fruit shelf life-related traits using two distinct F_2_ tomato populations. Fruit weight, firmness, shelf life (evaluated as both loss of fruit weight and loss of firmness over time), and colour space components were measured, and QTLs were mapped using a commercial low-density SNP genotyping panel and bulk segregant sequencing analysis experiments. We show that fruit firmness at harvest is only weakly predictive of postharvest firmness loss, indicating that shelf life should be treated as a dynamic trait rather than a static firmness phenotype. Across the two populations, 60 QTLs defining 26 genomic regions were identified, including both population-specific loci and shared regions on chromosomes 9 and 12. Several narrow intervals contained candidate genes related to ethylene signaling, cell-wall remodeling, calcium transport, aquaporin-mediated water balance and stress responses. The identified QTLs and candidate genes are directly relevant to breeding programmes seeking to improve postharvest performance without using major ripening mutants that compromise fruit quality.

**Key message:** Comparative QTL mapping and BSA-Seq in two F2 tomato populations show that postharvest shelf life is only partly explained by fruit firmness at harvest and identify candidate loci for firmness loss and weight loss during storage.

## Introduction

Tomato (*Solanum lycopersicum*) is one of the most widely cultivated vegetable crops worldwide and a major dietary source of vitamins and antioxidants. However, the postharvest shelf life of fresh market tomatoes remains a major limitation for supply chains, with fruit losses ranging from 20 to 50% in both developed and developing countries (Oladipo et al. 2026). Improving shelf life while maintaining fruit quality is therefore a major breeding objective.

The availability of genome and pangenome sequences (The Tomato Genome Consortium 2012; Gao et al. 2019), and large scale association studies for fruit quality traits (Zhao et al. 2019) have provided scientists and breeders with tools for studying genetic diversity and assisted breeding. Fruit quality traits such as the production of volatiles associated with flavor, primary metabolites and nutrient content have been mapped in tomato by Genome-Wide Association (GWA) approaches on whole populations and QTL mapping on segregating progenies (Çolak et al., 2023; Kaur et al., 2023 and references therein). GWAS and QTL analyses have revealed numerous loci controlling fruit quality traits (Kim et al. 2021; Zhang et al. 2024).

Tomato shelf life is determined by a complex combination of physiological and genetic factors, including fruit firmness, water loss, cuticle integrity, and the progression of ripening processes (Arah et al. 2015; Juan-Cabot et al. 2022). Fruit softening is primarily driven by cell wall remodeling enzymes such as polygalacturonases, pectin methyl esterases and expansins, whereas water loss is strongly influenced by cuticle composition and epidermal properties. Hormonal regulation also plays an important role, with ethylene signaling and abscisic acid metabolism influencing ripening progression and postharvest physiology (Bassolino et al. 2013; Zhang et al. 2018; Diretto et al. 2020; Fernández-Muñoz et al. 2022; Mubarok et al. 2023). Unlike fruit firmness measured at harvest, shelf life reflects the cumulative outcome of multiple physiological processes occurring during postharvest storage. Although fruit firmness at harvest is widely used as a proxy for shelf life in tomato breeding, it remains unclear to what extent these traits share the same physiological and genetic basis.

Since the early demonstration of the role of polygalacturonase in tomato ripening through antisense RNA (Oeller et al. 1991), several genes affecting tomato fruit firmness and shelf life have been identified. Classical ripening mutants such as *rin*, *nor*, and *Cnr* strongly affect fruit softening but also compromise fruit sensory quality (Wang et al. 2020). The role of *Alc*, another allele of *NOR* gene, has been also demonstrated (Yogendra and Ramanjini Gowda 2013). The *FIS1* locus, encoding a GA2-oxidase enzyme, was shown to modulate fruit firmness by altering pericarp cell expansion (Li et al. 2020). The role of other genes has been shown by gene editing of a pectate lyase (Fumelli et al. 2026), a histone deacetylase *SlHDA7* (Zhou et al. 2024), a Lateral Organ Boundaries transcription factor, *SlLOB1* (Shi et al. 2021), and simultaneous knockout of two pectin-degrading polygalacturonase and pectate lyase enzymes (Ortega_-_Salazar et al. 2024).

Despite these advances, the genetic architecture of postharvest shelf life remains poorly understood compared with other fruit quality traits. Shelf life is a complex quantitative trait influenced by multiple processes including water loss, softening rate and ripening dynamics. A few QTL studies have focused on fruit firmness on single populations (Causse 2002; Diouf et al. 2018) or GWAS panels (Albert et al. 2016; Kayikci et al. 2025). A genomic region involved in fruit firmness was also studied and dissected into several QTLs with epistatic interactions (Chapman et al. 2012).

Nevertheless, the relationship between shelf life and fruit firmness QTLs remains unclear and there are only a few studies on the genetic control of shelf life in natural diversity. The analysis of a *S. pennellii* x *S. lycopersicum* introgression line population allowed the identification of several regions and suggested the role of a few candidate genes for shelf life performance (Uluisik and Oney-Birol 2021). Several genomic regions involved in shelf life variation were also identified in a RIL population derived from an interspecific *S. lycopersicum* x *S. pimpinellifolium* cross (Cambiaso et al. 2019). To our knowledge, studies investigating the genetic basis of fruit shelf life in intraspecific *S. lycopersicum* segregating populations remain very limited.

Comparative mapping across multiple populations provides a powerful approach to identify both conserved and population-specific loci controlling complex traits. Furthermore, combining classical QTL mapping with bulk segregant sequencing analysis (BSA-Seq) can improve the detection and refinement of genomic regions associated with extreme phenotypes (Majeed et al. 2022).

In the present study, we investigated the genetic basis of tomato fruit shelf life using two intraspecific F2 populations derived from independent crosses. Fruit weight, firmness, and color traits were monitored during postharvest storage, allowing the quantification of both initial fruit characteristics and their evolution during storage. QTL mapping based on a SNP genotyping panel was complemented with BSA-Seq analysis of extreme phenotypes. This combined approach was used to identify genomic regions associated with fruit firmness, weight loss and other quality traits, revealing both shared and population-specific loci contributing to tomato shelf life. As parental genomes were re-sequenced, polymorphisms in candidate genes were screened and analysed in relation to their potential impact on trait expression.

## Materials and Methods

### Plant material and growth and phenotyping procedures

The two tomato (*S. lycopersicum*) F2 populations used in this study had different origins and pedigree. The parental lines of ENEA F2 population (G2P-SOL codes: P1 GPT005160; P2 GPT026210) were selected from the G2P-SOL (https://www.g2p-sol.eu/) core collection for contrasting fruit firmness and an average fruit mass within the second quantile of the whole collection (41.3-77.2 g range).

The INRAE F2 population was derived by selfing the F1 commercial hybrid Garance bred by INRAE (https://eng-gafl.paca.hub.inrae.fr/results/plant-breeding-innovation/varietal-vegetable-creation), characterized by large fruit mass, high firmness and long shelf life.

In spring-summer 2023, both F2 populations were soil-grown at INRAE Avignon in the same greenhouse. The F2 plants showing determinate growth habit were discarded. ENEA F2 progenies were grown together with P1, P2 and F1 plants, each grown in eight random replicates. A total of 215 ENEA F2 progenies were selected for phenotyping analyses. INRAE F2 progenies were grown together with Garance F1 and both parents, each grown in ten random replicates. A total of 219 INRAE F2 progenies were selected for phenotyping analyses. Growth conditions were as previously described (Pascual et al. 2015).

### Phenotyping procedures

Fruits were harvested at the breaker stage, and the following traits were measured on fruits (five replicates per plant) harvested from trusses 2 to 6 (**Tables S1** and **S2**):

- fruit weight (FW) at harvest (FW_T0_), 7 days (FW_T7_) and 14 days after harvest (FW_T14_);
- fruit firmness (FF), measured at two opposite equatorial positions at T0 (FF_T0_) and T14 (FF_T14_) with a 5-mm diameter flat tip durometer (TR©Turoni, ENEA; Durofel, INRAE, daily calibrated the same way).
- fruit color space components (L*, brightness; a*, red/green component, and b*, yellow/blue component) measured at two equatorial opposite positions at T0 (L*_T0_, a*_T0_, b*_T0_) and T14 (L*_T14_, a*_T14_, b*_T14_) with a CR400 colorimeter (Konica Minolta).

Following T0 measurements, fruits were kept in the dark at 18 °C until measurements at T7 and T14, to simulate postharvest handling conditions. Shelf life was assessed during storage with two parameters: the loss of FW at T7 and T14, compared to FW_T0_ (LossFW_T7_ = 1 - FW_T7_/FW_T0_; LossFW_T14_ = 1 - FW_T14_/FW_T0_) and the loss of FF at T14 with respect to FF_T0_ (LossFF_T14_ = 1 - FF_T14_/FF_T0_). Variations of L*, a* and b* during storage were calculated as the difference between values at T14 and T0, to obtain delta L*_T14_, delta a*_T14_ and delta b*_T14_.

### Genotyping and QTL mapping for fruit quality traits

Lyophilized leaf samples (two 6-mm leaf discs per genotype) were sent to Agriplex Genomics (Cleveland, OH, USA) for low-density genotyping using their 1078-SNP Commercial Tomato Panel (https://www.agriplexgenomics.com/products/public-panels/commercial-tomato-panel). Genotype data are presented in **Tables S3** and **S4**. A genetic map was then constructed using the R/qtl package (Broman et al. 2003), following authors’ instructions. Phenotypic data from each population were analysed independently following the same workflow (test of normality of the distribution, average calculation per F2 plant, correlations among traits) using R software (version 3.6.2). QTLs were mapped on traits, transformed when necessary to reach normality, using Composite Interval Mapping (CIM) model for either population using the R/qtl package. The LOD threshold was calculated for each trait following 1000 permutations and a 5% genome-wide significance level.

### Bulk segregant sequencing analysis (BSA-Seq)

Twelve to 19 extreme F2 individuals for selected shelf life traits were used in each bulk (selected traits: FF_T14_, LossFW_T14_ for ENEA; FF_T0_ and LossFF_T14_ for INRAE; **Tables S1 and S2**). Genomic DNA was isolated using the DNeasy Plant Mini Kit (Qiagen) from 10-mg samples (eleven 8-mm disc punches) of lyophilized young leaves. DNA quality control was performed by agarose gel electrophoresis to evaluate RNA contamination and DNA integrity. DNA concentration was determined using a fluorimetric method based on Quant-iT™ PicoGreen™ (ThermoFisher Scientific) and a microplate reader (Varioskan LUX, ThermoFisher Scientific). The bulked DNA samples were prepared by mixing equal amounts of DNA (200 ng for each genotype) extracted from each segregant pool. Bulk DNA samples were shipped to an external provider (BGI) for library preparation (short-insert size library, PCR based) and paired-end 150 bp resequencing at 40x coverage on a MGI/DNBSEQ platform. The fastq files were quality-filtered using fastp (Chen 2023) with default options and QTL-seq package (Sugihara et al. 2022) was used for BSA-Seq analysis. The *S. lycopersicum* genome v.4.00 was used as reference and default parameters were used except for an additional filtering step of the vcf file obtained by BSA-Seq. This consisted, for both biparental populations, in extracting only those biallelic loci on the genome where the two parental accessions had homozygous contrasting alleles.

### Search for shelf life candidate genes

The SL4.0 Tomato Genome version was used to select candidate genes for shelf life QTLs (FF, LossFF, FW, and LossFW traits). QTLs derived from CIM and BSA-Seq with confidence intervals <2.5 Mb were screened for genes with functional relevance to the traits, including gene ontology analysis. Candidate genes were prioritized based on their annotated biological function and their putative involvement in fruit firmness, fruit weight and postharvest physiology. Their expression profiles during fruit development and ripening were then examined using the Tomato Expression Atlas (https://tea.solgenomics.net/; Fernandez-Pozo et al. 2017). Genes showing no detectable expression in fruit developmental or ripening stages were excluded from further analyses, whereas genes for which expression data were unavailable were retained and labelled as NA. The four parental lines were sequenced and polymorphisms in these genes between pairs of parents were identified following the procedures presented by Desaint et al. (2024). Variant position and predicted functional impact were determined using ANNOVAR (Wang et al. 2010).

## Results and Discussion

Two F2 segregating populations were used in this study: (1) the progeny from an F1 commercial hybrid called Garance, selected at INRAE, for its good fruit quality and shelf life, despite the absence of *rin* or *nor* mutations, and (2) the progeny derived from a cross between two parental lines selected from the G2PSOL tomato core collection, contrasting for fruit firmness. These populations will be hereinafter referred to as INRAE and ENEA populations.

### Phenotypic diversity of the two populations

The main phenotypic characteristics of the parent accessions and F2 populations are shown in **Table 1**. The fruit weight (FW) at the fully ripe stage of the F2 progenies ranged between 36.7 and 224.8 g (ENEA population) and between 30.5 and 166.3 g (INRAE population) (**Figure 1A-B**). In both populations, there was a significant difference in FW between parental lines. Fruits of both F1 hybrids had opposite FW values: FW of ENEA F1 was almost identical to the small-fruited parent, while INRAE F1 had a FW closer to the large-fruited parent (**Figure 1A-B**, **Tables 1** and **S1-S2**), suggesting a different genetic control. FW had high repeatability. Fruit weight decreased with a linear trend during the 14-day storage period and distributions for FW showed the same pattern at T7 and T14 (data not shown). LossFW_T14_ in ENEA F2 population ranged from 1.5% to 6.1% (average of F2 population: 2.7%) (**Table 1**, **Figure 1C**). The F1 hybrid lost more FW than the parental lines throughout the 14-day storage (**Table 1**, **Figure 1C**). On the other hand, LossFW_T14_ in INRAE F2 population had a wider variation range (1.4-10.1%), with an average value close to that of P1 and F1 genotypes (**Table 1**, **Figure 1D**). Repeatability of LossFW_T14_ was lower than FW in both populations.

**Figure 1.**
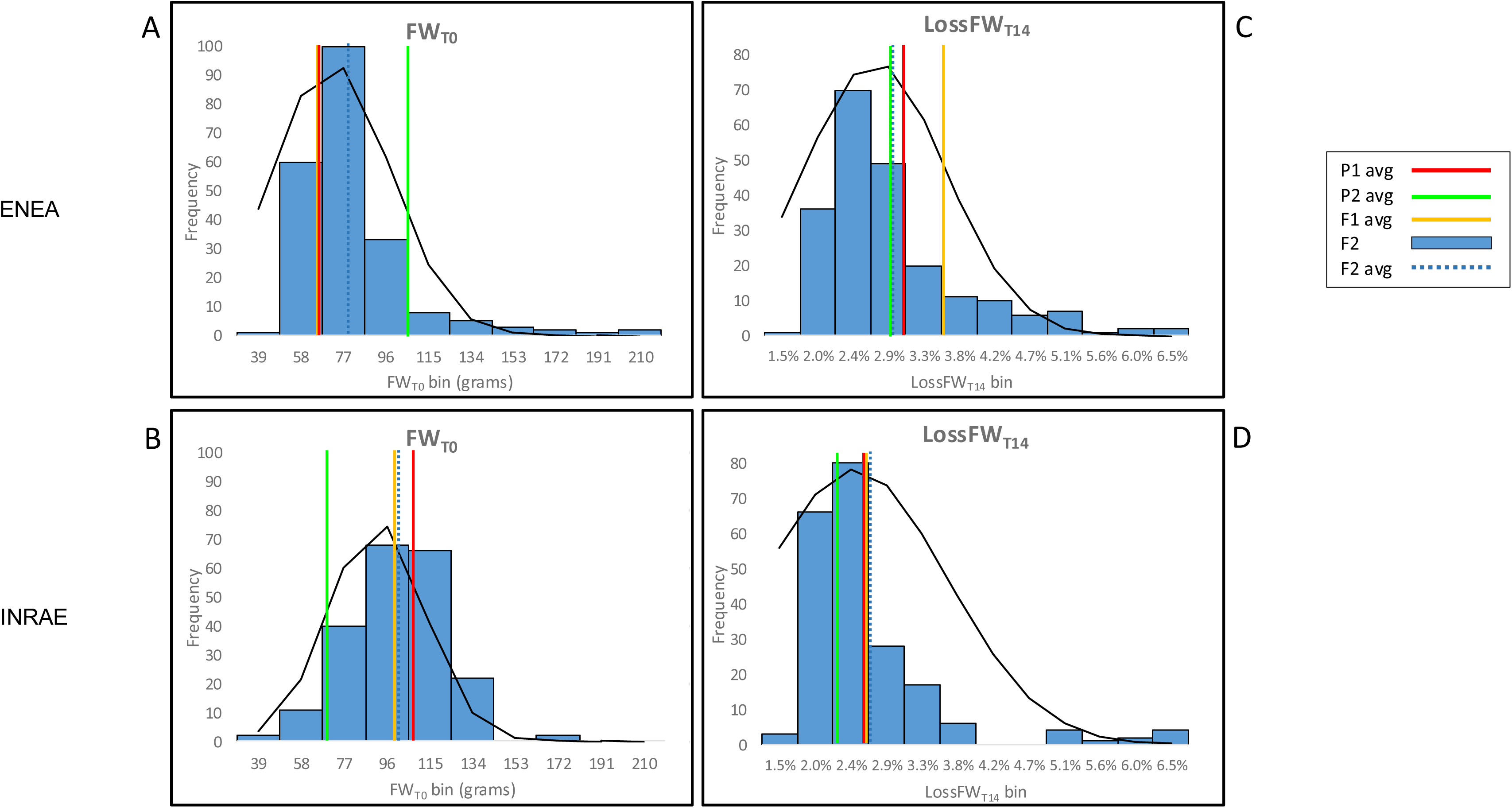
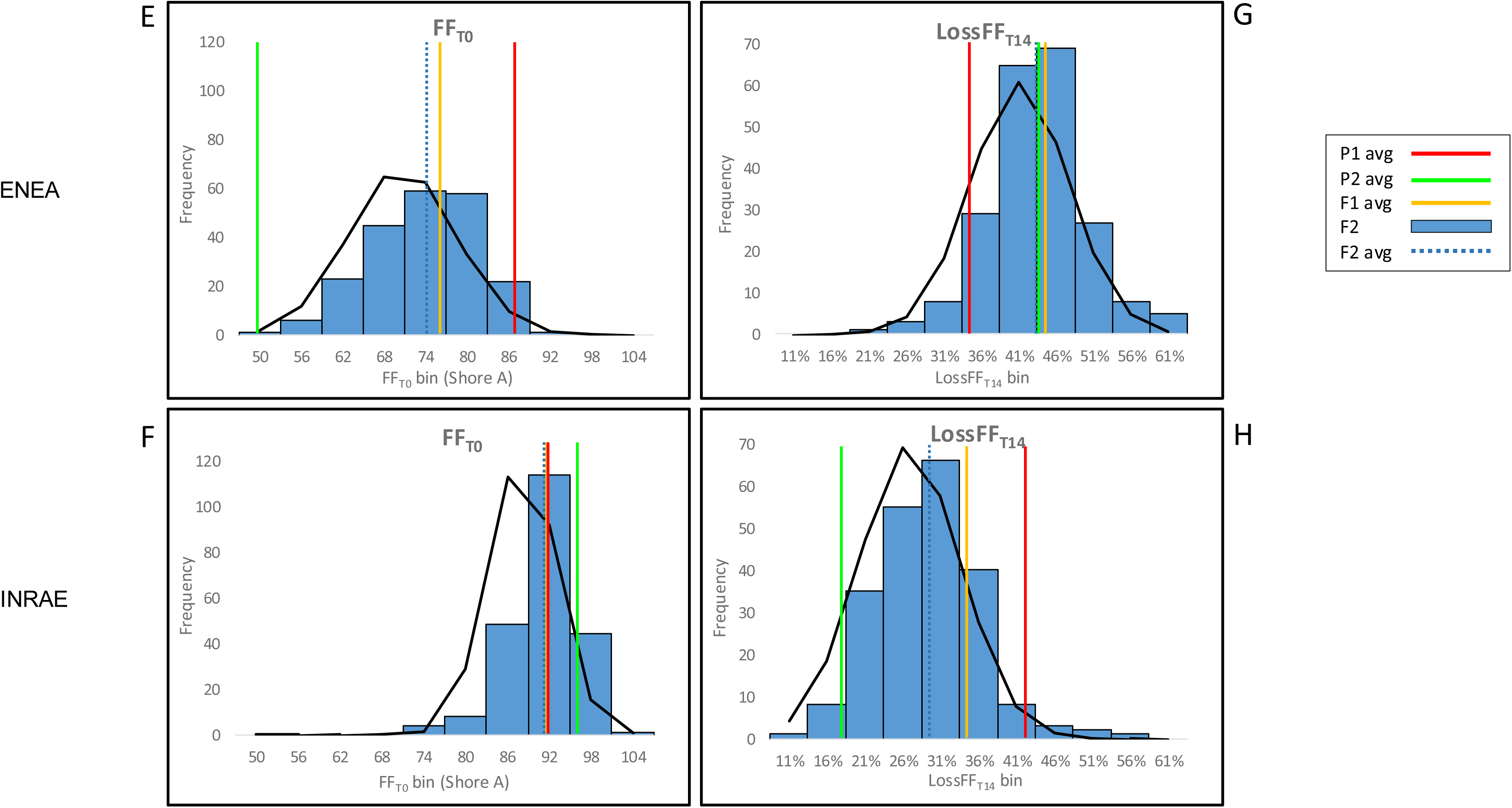
Distribution of FW- and FF-related traits in studied F2 populations. A-B: Distribution of FW_T0_ in ENEA (A) and INRAE (B). C-D: Distribution of LossFW_T14_ in ENEA (C) and INRAE (D). E-F: Distribution charts of FF_T0_ in the ENEA (E) and INRAE (F). G-H: Distribution of LossFF_T14_ in the ENEA (G) and INRAE (H). Blue histograms: F2 progeny bins. Dark blue dotted line: average of F2 population. Red line: average of P1. Green line: average of P2. Orange line: average of F1 hybrid. Black line: distribution curve of F2 population.

**Table 1.** Summary of the characteristics of the parental lines, their F1 hybrids and the two F2 populations.

| Trait | ENEA population |  |  |  |  |  |  |  | INRAE population |  |  |  |  |  |  |  |
| --- | --- | --- | --- | --- | --- | --- | --- | --- | --- | --- | --- | --- | --- | --- | --- | --- |
|  | P1<br>avg | P2<br>avg | F1<br>avg | F2<br>min | F2<br>max | F2<br>avg | F2<br>sd | var | P1<br>avg | P2<br>avg | F1<br>avg | F2<br>min | F2<br>max | F2<br>avg | F2<br>sd | var |
| FW<sub>T0</sub> | 57.4 | 97.0 | 56.7 | 37.7 | 229.9 | 71.7 | 26.3 | 0.47 | 98.3 | 60.6 | 90.4 | 33.8 | 169.7 | 91.5 | 21.0 | 0.56 |
| FW<sub>T7</sub> | 56.4 | 95.5 | 55.6 | 37.1 | 226.9 | 70.6 | 26.1 | 0.47 | 96.7 | 59.9 | 89.2 | 30.9 | 168.0 | 90.2 | 20.9 | 0.56 |
| FW<sub>T14</sub> | 55.9 | 94.4 | 54.9 | 36.7 | 224.8 | 69.9 | 25.9 | 0.47 | 96.0 | 59.3 | 88.3 | 30.5 | 166.3 | 89.3 | 20.7 | 0.56 |
| LossFW<sub>T7</sub> | 1.8 | 1.5 | 2.0 | 0.8 | 4.1 | 1.6 | 0.6 | 0.31 | 1.6 | 1.0 | 1.5 | 0.6 | 8.7 | 1.4 | 1.0 | 0.20 |
| LossFW<sub>T14</sub> | 2.8 | 2.7 | 3.4 | 1.5 | 6.1 | 2.7 | 0.9 | 0.31 | 2.4 | 2.1 | 2.4 | 1.4 | 10.1 | 2.5 | 1.2 | 0.20 |
| FF<sub>T0</sub> | 83.8 | 46.6 | 73.1 | 48.4 | 86.8 | 70.7 | 7.8 | 0.43 | 88.4 | 93.0 | 88.4 | 69.3 | 98.8 | 88.2 | 4.8 | 0.44 |
| FF<sub>T14</sub> | 56.9 | 27.2 | 42.1 | 29.2 | 57.5 | 41.4 | 5.3 | 0.48 | 53.2 | 78.8 | 60.3 | 39.9 | 80.6 | 64.5 | 7.3 | 0.54 |
| LossFF<sub>T14</sub> | 32.0 | 41.4 | 42.3 | 17.3 | 60.4 | 41.2 | 6.5 | 0.21 | 39.7 | 15.2 | 31.8 | 8.9 | 54.9 | 26.9 | 6.7 | 0.35 |
| L*<sub>T0</sub> | 56.8 | 53.7 | 54.5 | 47.9 | 65.5 | 54.7 | 2.6 | 0.26 | 58.6 | 54.8 | 54.2 | 45.9 | 61.0 | 53.5 | 2.4 | 0.28 |
| a*<sub>T0</sub> | 6.0 | 10.4 | 8.4 | -5.8 | 24.1 | 5.9 | 4.7 | 0.13 | 5.1 | -0.19 | 0.2 | -6.29 | 24.5 | 2.6 | 5.1 | 0.01 |
| b*<sub>T0</sub> | 33.3 | 19.7 | 29.3 | 20.0 | 38.0 | 29.9 | 4.3 | 0.66 | 23.2 | 36.3 | 27.7 | 1.2 | 40.9 | 28.4 | 6.5 | 0.11 |
| L*<sub>T14</sub> | 41.7 | 41.1 | 39.7 | 37.8 | 45.2 | 40.5 | 1.3 | 0.55 | 42.1 | 42.8 | 40.6 | 35.3 | 45.4 | 40.7 | 1.5 | 0.48 |
| a*<sub>T14</sub> | 32.9 | 24.0 | 28.7 | 22.5 | 35.5 | 28.7 | 2.4 | 0.72 | 29.2 | 28.7 | 26.6 | 16.9 | 33.0 | 27.3 | 2.2 | 0.42 |
| b*<sub>T14</sub> | 29.3 | 16.8 | 24.2 | 15.2 | 33.1 | 24.0 | 3.7 | 0.82 | 30.1 | 32.1 | 26.9 | 19.2 | 35.9 | 28.6 | 2.6 | 0.43 |
| delta L*<sub>T14</sub> | -15.0 | -12.6 | -14.8 | -21.3 | -8.3 | -14.2 | 2.1 | 0.55 | -16.4 | -12.0 | -13.6 | -18.4 | -6.5 | -12.8 | 2.1 | 0.20 |
| delta a*<sub>T14</sub> | 26.9 | 13.6 | 20.3 | 4.7 | 37.3 | 22.8 | 5.1 | 0.18 | 24.1 | 28.9 | 26.4 | 5.0 | 34.6 | 24.7 | 4.9 | 0.01 |
| delta b*<sub>T14</sub> | -4.0 | -2.9 | -5.1 | -12.4 | 1.3 | -6.0 | 2.7 | 0.36 | 6.9 | -4.1 | -0.8 | -12.0 | 22.9 | 0.3 | 6.3 | 0.05 |

Avg: Average values from 3 harvests of 5 fruits/plant (see Methods). Sd: standard deviation; var: repeatability across the three harvests

The two populations exhibited striking differences for fruit firmness (FF). P1 and P2 from the ENEA population were strongly contrasted with FF, P1 fruits being significantly firmer than those of P2 at either T0 or T14, while F1 fruits had intermediate values. ENEA F2 population showed a large variation of FF, ranging between 48.4 and 86.8 shoreA degrees (average of F2 population: 70.7) at harvest (**Tables 1** and **S1**) with a normal distribution pattern (**Figure 1E**). In contrast, the distribution pattern of INRAE F2 population was much narrower and shifted towards values higher than in the ENEA population (69.3-98.8 range, 88.2 on average) (**Tables 1** and **S2; Figure 1F**). This probably reflects the origin of the INRAE population from a commercial hybrid selected for high fruit firmness. At the end of storage, FF_T14_ markedly decreased in both populations (ENEA: 29.2-57.5 range; INRAE: 39.9-80.6 range). While most F2 progenies of the ENEA population had FF_T14_ falling within parental range, 16 out of 219 F2 plants exhibited FF lower than the softer parental in the INRAE F2 population. The ENEA F2 population had a wide variation of LossFF_T14_, from 17.3% to 60.4% (**Table 1**, **Figure 1G**). Fruits of the firmest parent P1 had a significantly lower LossFF_T14_ than those of P2 and F1 hybrid (**Figure 1G**). The origin of a firm, commercial hybrid of INRAE population caused the observed shift in the LossFF_T14_ towards lower values: LossFF_T14_ values of F2 progenies ranged between 8.9% to 54.9%, with Garance F1 at 31.8%, a value intermediate between those of F2 average and P1 (**Figure 1H**). The repeatability of FF and LossFF was close to those of FW and Loss FW. Overall, the two populations differed markedly in both initial fruit firmness and firmness retention during storage, providing complementary genetic backgrounds for investigating the genetic basis of tomato shelf life.

Fruit skin brightness **L\*** had similar ranges of variation at harvest (T0) in both populations (**Figure S1**, **A** and **B**) and decreased during storage (delta L*_T14_ negative values) (**Table 1**). The possible reasons for this decrease could be accumulation of pigments such as carotenoids/flavonoids, and thus an increase in absorbance, or fruit softening and dehydration, which led to a decrease of surface reflectance (Vicente et al. 2007; Kapoor et al. 2022). Compared to parental lines, ENEA F1 fruits showed intermediate values for both L*_T0_ and delta L*_T14_ (**Figure S1**, **A** and **C**). ENEA P1 fruits had higher L*_T0_ than P2 ones, and lower delta L* during storage (**Figure S1C**). The INRAE F2 population was characterized by higher median values (**Figure S1B**). The INRAE P1 parent had significantly higher L* values than P2, F1 and F2 average, but showed a stronger decrease in skin brightness during storage (**Figure S1D**).

The **a\*** parameter describes the green-to-red axis of the color space. It showed a strong increase from T0 (harvest at breaker stage) to T14 (red ripe stage), consistent with the accumulation of lycopene during ripening (**Table 1, Figure S2**). During storage, ENEA P1 fruits had an increase in a* values significantly higher than P2 and F1, with F1 hybrid showing a* and delta a* intermediate values halfway between parental lines (**Table 1, Figure S2**, **A** and **C**). INRAE F2 population showed lower a* values at harvest (**Figure S2**, **B** and **D**), which were compensated by a distribution frequency towards higher delta a*_T14_ values at the end of storage (**Figure S2D**).

The **b\*** component describes the blue-to-yellow axis of the color space. The lower b*_T0_ values of the ENEA P2 parent (**Figure S3A**) are consistent with its *pink* phenotype, impairing the accumulation of yellow pigments in fruit skin (Ballester et al. 2009). The ENEA F1 has **b\*** values similar to the P1 parent, in agreement with the recessive nature of the *pink* mutation. During storage, b* values generally decrease during storage, in agreement with the decrease in flavonoids (Carrari 2006; Del Giudice et al. 2015) with the exception of the INRAE P1 parent, which shows a slight increase (**Table 1; Figure S3C**). This is reflected in the wider distribution of both b*_T0_ and delta b*_T14_ values in the INRAE F2 population (**Figure S3**, **B** and **D**). The repeatability in these components was high, with the exception of a* at T0 in INRAE population where the color of harvested fruits was highly variable. It was much more repeatable 14 days after harvest. Overall, the observed variation in color traits was consistent with normal fruit ripening during postharvest storage and highlighted the broad phenotypic diversity of the two F2 populations.

### Correlation among traits

Correlation analyses were carried out to unveil possible associations between the traits in each F2 population (**Figure 2**). Strong positive correlations were found between FW or LossFW traits measured at different time points, indicating that water loss progressed in similar linear patterns in the different plants of the F2 progenies. The negative correlations between L*_T14_ and LossFW indicate the role of skin structure and composition in regulating water loss and hence fruit weight and freshness (Ji et al. 2023). The weak negative correlations between FW and LossFW (average ENEA −0.31, INRAE −0.25) were expected, since heavier fruits had a larger volume/surface ratio and hence lost water more slowly. Similarly, the negative correlations between trait pairs FF_T14_-LossFF_T14_, L*_T0_-delta L*_T14_, a*_T0_-delta a*_T14_, and b*_T0_-delta b*_T14_ were also expected, since the second trait is derived from the first. Likewise, positive correlations between pairs b*_T0_-b*_T14_, and L*_T0_ with other color traits L*_T14_, a*_T14_ and b*_T14_ were observed in both populations. Notably, the low correlations between LossFW and either FF or LossFF traits (|*r*| ≤ 0.28) indicate the lack of relationship between these two main determinants of shelf life. Such absence or low correlations between FW and FF were also found in other populations and point out the different genetic control of these complex traits (Saliba-Colombani et al. 2001; Albert et al. 2016; Diouf et al. 2018). Among the correlations found only in a single population, the negative correlation between a*_T0_ and FF_T0_ (−0.59) in ENEA population can be explained by the fact that fruits that are slightly earlier in the ripening curve are expected to contain less lycopene and be firmer.

**Figure 2.**
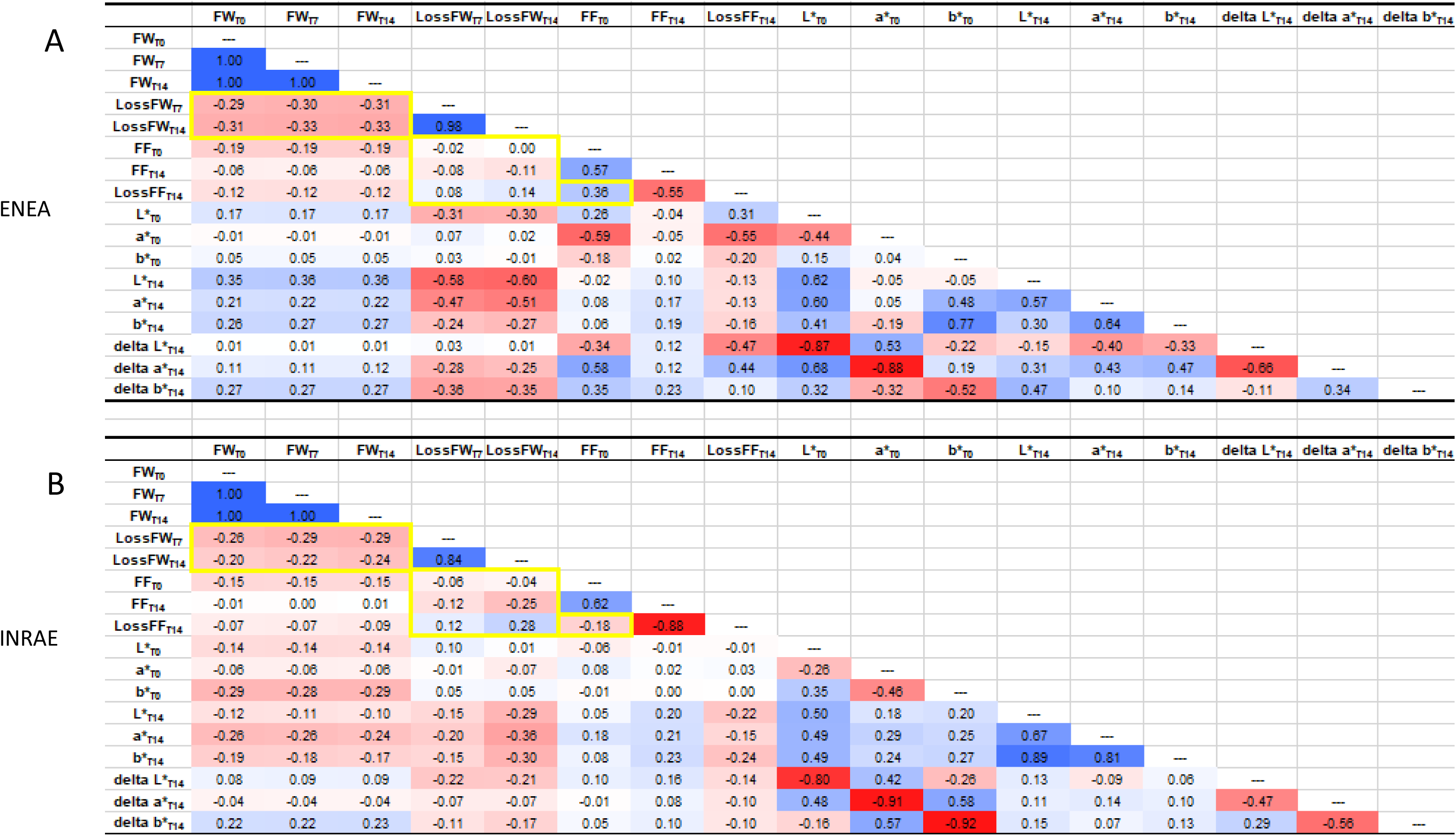
Pearson correlation matrices between traits in studied F2 populations. A: ENEA. B: INRAE. a*: red/green color; b*: yellow/blue color; delta a*_T14_: a*_T14_ - a*_T0_; delta b*_T14_: b*_T14_ - b*_T0_; delta L*_T14_: L*_T14_ - L*_T0_; FF: fruit firmness; FW: fruit weight; L*: fruit skin brightness; LossFF_T14_: loss of FF at T14; LossFW_Tx_: loss of FW at T7 or T14. Blue: positive correlation. Red: negative correlation. White: no correlation.

Although fruit firmness at harvest (FF_T0_) is widely used as a proxy for tomato shelf life, it was only weakly correlated with firmness loss during storage (LossFF_T14_) in both populations. **Figure S4** illustrates the different patterns of relationships for traits related to shelf life in both populations. A more linear relationship between FF_T14_ and LossFF_T14_ was observed in the INRAE population than in ENEA one, whereas the triangular relationship between FW_T0_ and LossFW_T14_ or LossFW_T14_ and LossFF_T14_ detected in the ENEA population was absent in the INRAE population, suggesting distinct genetic control of the traits. Overall, the correlation analyses indicate that the major components contributing to tomato shelf life are only partially associated, supporting the hypothesis that they are, at least in part, under distinct genetic control.

### SNP diversity and genetic maps

The two populations were genotyped with the Agriplex Fresh Market Tomato 1K panel (https://www.agriplexgenomics.com/), showing 27% and 24% polymorphism and uneven distribution of SNPs in both populations (**Figure S5A**). This can be related to the different distribution of SNPs in the genomic sequences of the parental lines (**Figure S5**, **B** and **C**). The construction of genetic maps of both populations revealed some clusters of markers corresponding to highly polymorphic regions and to regions with low recombination (**Figures S6A** and **S6B**). Both population maps highlighted a cluster of polymorphic markers in the short arm of chr. 2 (0-30 Mb), corresponding to the acrocentric region of this chromosome (Sim et al. 2012). A high density of polymorphic markers was also observed in the distal region of the long arm of chromosome 2, corresponding to a hotspot for agronomically important QTLs and genes as *dblk2.1*, *fw2.2*, *lc*, *ovate*, regulating fruit shape and size, locule number, and plant development (Alpert et al., 1995; Frary et al., 2000; Liu et al., 2002; Ranc et al., 2012) (**Figures S6A** and **S6B**). This uneven distribution of polymorphisms led to some chromosomes very poorly saturated (chr. 4 of ENEA map; chr. 1 and 4 of INRAE map) or chromosomes separated in two linkage groups (chr. 6, 10, 11 of ENEA map and chr. 6, 9, 12 of INRAE map). This is likely to be responsible for the low number of QTLs detected in these chromosomes. The ENEA map included additional polymorphic marker clusters on chr. 8, 9, 11 and 12 (**Figure S6A**), while the INRAE map contained marker clusters on chr. 3 and 10 (**Figure S6B**). Overall, the uneven distribution of polymorphic markers resulted in variable map coverage among chromosomes, a factor that should be considered when comparing the number and location of detected QTLs.

### Multi-approach mapping for fruit quality- and shelf life-related QTLs

To identify genomic regions controlling fruit quality and postharvest shelf life, QTLs were first mapped through composite interval mapping (CIM). To complement this approach and capture additional loci associated with shelf life performance, two BSA-Seq experiments were carried out by selecting extreme segregants for FF_T14_ and LossFW_T14_ (ENEA) and FF_T0_ and LossFF_T14_ (INRAE). **Figure 3** summarizes the QTL intervals for FW- and FF-related traits obtained by both mapping approaches (CIM and BSA-Seq).

**Figure 3.**
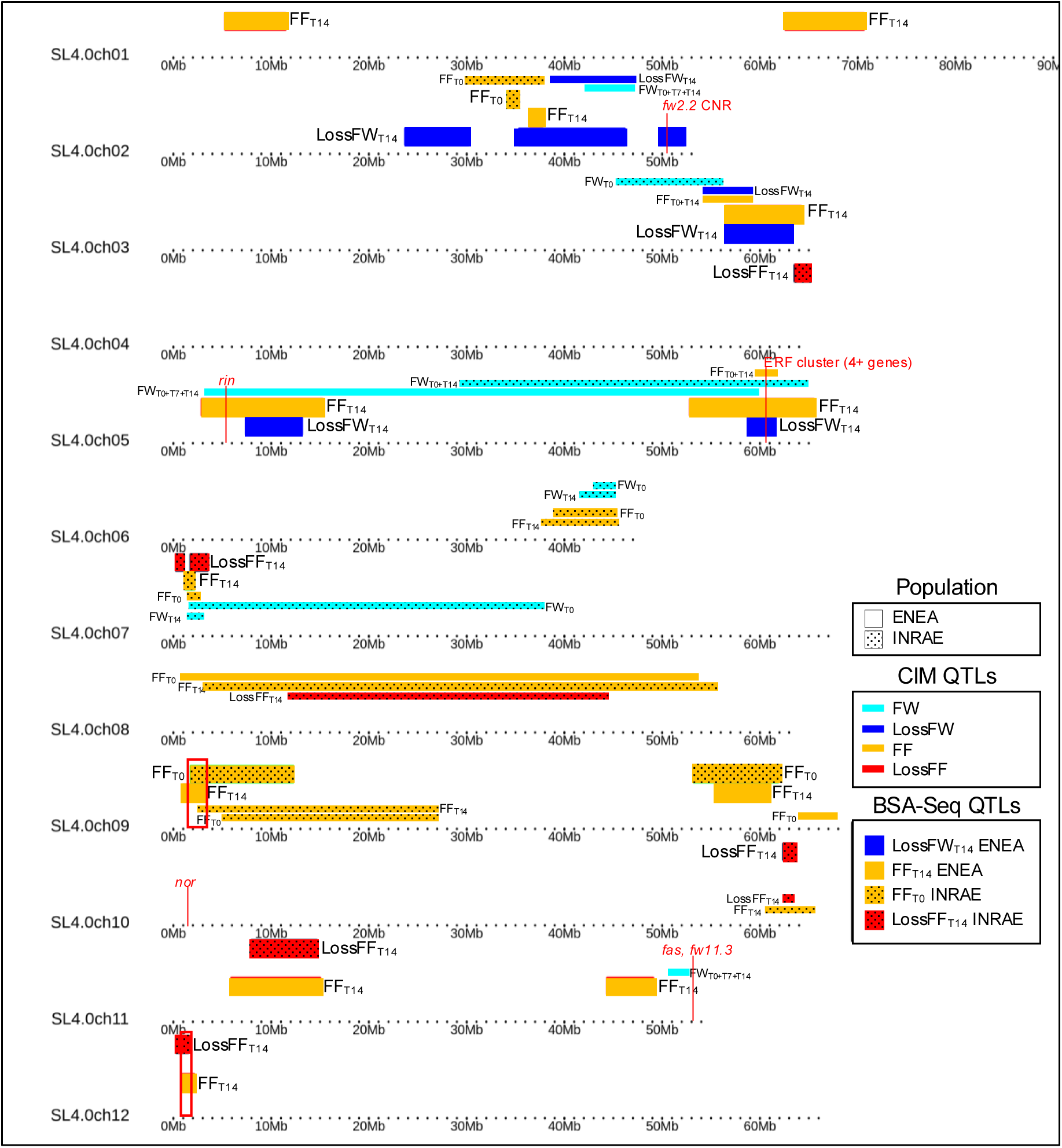
Combined QTL map for shelf life-related FW and FF traits. QTLs from genetic maps (detected by CIM) and BSA-Seq of both ENEA and INRAE F2 populations are shown. Key ripening-related genes in QTL regions were added as reference (in red).

#### CIM for fruit quality traits

Genetic maps of QTLs for FW, FF and fruit color space components as well as for derived traits following fruit storage detected by CIM for each population are shown in **Figures S7 A** and **B**. The main QTL features are shown in **Table 2** and more detailed information in **Tables S5** and **S6**. In the ENEA population, 13 QTLs were detected by CIM for FW- and LossFW-related traits on chr. 2, 3, 5 and 11B (**Tables 2** and **S5**). The QTLs on chr. 2 could be associated with *fw2.2*, located in the bottom part of chr. 2 and regulating fruit size and weight (Alpert et al. 1995; Frary et al. 2000). Nevertheless the large fruit allele of FW2.2 has been fixed after domestication and several QTLs for FW have been detected in the region by fine mapping (Lecomte et al. 2004). On chr. 3, only a QTL specific for LossFW_T14_ was found. The FW-associated QTL on chr. 11B had a very strong effect (LOD = 19). It was close to the *fasciated* and the *fw11.3* loci, known to control fruit shape and size, respectively (Huang and Van Der Knaap, 2011; Rodríguez et al., 2011; Mu et al, 2017). All these QTLs were mainly additive, with the same parent providing the strong effect allele, except for LossFW _T14_, where the two QTLs had opposite effects. For FF, QTLs were detected on chr. 3, 5 (with a strong effect) and 8. QTLs for FF_T0_ and FF_T14_ overlapped on chr. 3 and 5, the FF_T0_-associated QTLs having a smaller confidence interval. The QTLs for LossFW_T7_ and LossFW_T14_ co-localized with those for FF_T0_ and FF_T14_ on chr. 3. Surprisingly, no QTLs were detected for LossFF_T14_. QTLs associated with color space components L* (fruit skin brightness), a* (red/green color) and b* (yellow/blue color) were mapped on several chromosomes in the ENEA population. L*-related QTLs were located on chr. 2, 3, 5, 8 and 11B, with chr. 3 and 5 containing overlapping QTLs for L*_T0_ and L*_T14_ (in both cases, L*_T0_ QTLs had a smaller confidence interval). QTLs for a* traits mapped on chr. 1, 3, 5, 8 and 9. The only co-mapping QTLs for a*_T0_ and a*_T14_ were present on chr. 1, while strong a*_T14_ QTLs were located on chr. 3 and 8. QTLs for b* traits were present on chr. 1, 2, 5, 8, 10 and 11. Overlapping QTLs for b* _T0_ and b* _T14_ were detected on chr. 1 and 5. The former QTL was the strongest QTL for the ENEA population (chr. 1, 65.1-73.1 Mb), accounting for over 34% of the genetic variation for b* traits (**Table S5**), and corresponded to the region carrying the *y* mutation in a *Myb12* gene (*Solyc01g079620*) conferring the colorless epidermis and *pink* fruit phenotype (Ballester et al. 2009). QTLs for all color components co-mapped on chr. 5 and 8: in the first case, they overlapped also with FW and FF_T0_ QTLs, while on chr. 8 they co-localised only with a FF_T0_ QTL (**Figure S7A**).

**Table 2.**
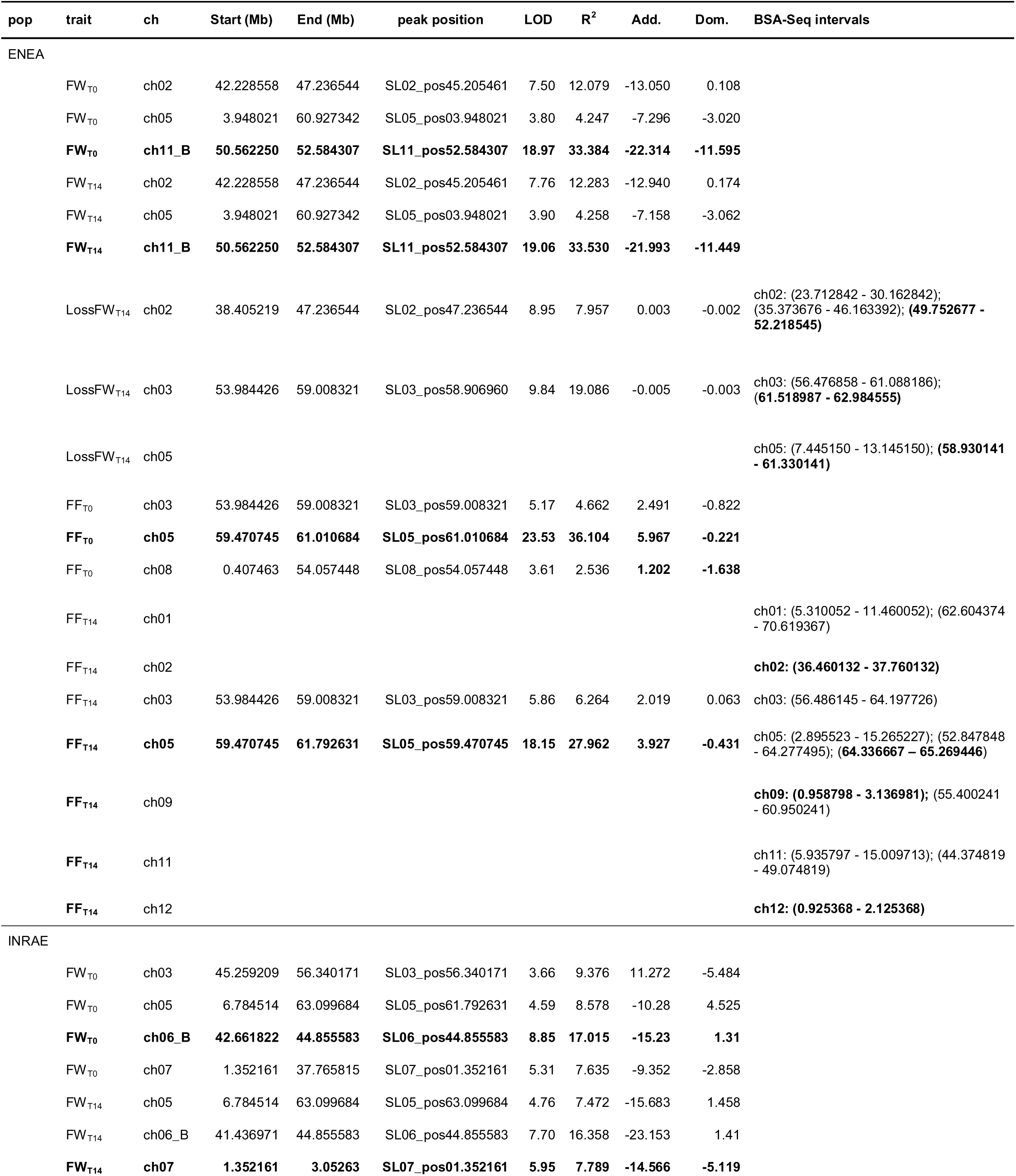
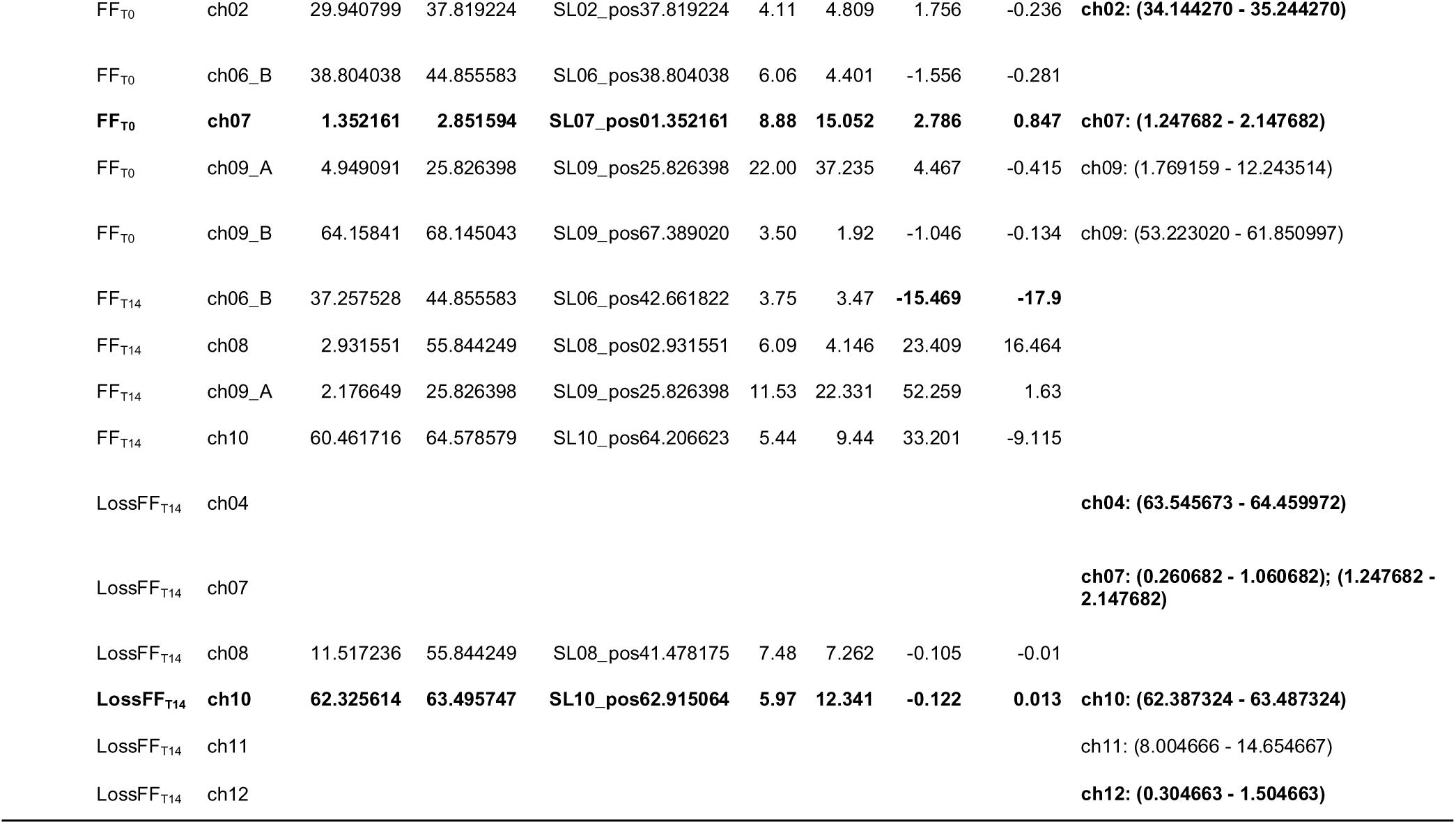
QTL information in the two genetic maps for FW, LossFW, FF and LossFF traits at T0 and T14, obtained by Composite Interval Mapping and BSA-Seq approaches. QTLs with Confidence Interval <2.5 Mb, further searched for candidate genes, are highlighted in bold.

In the INRAE population, FW-related QTLs were detected on chr. 3, 5, 6B and 7 at T0, and in the same regions except on chr. 3 at T14, but no QTLs were detected for LossFW_T14_ (**Tables 2** and **S7B**). The QTL on chr. 6B had the highest LOD score and explained more than 17% of the variation. It could correspond to the *fw6.3* QTL which was located in the *sp* gene region, controlling determinate growth (Ning et al. 2023). Determinate plants homozygous for *sp* were discarded, but the contrast between heterozygous and homozygous *sp+* remained for analyses. QTLs for FF-related traits were located on chr. 2, 6B, 7, 8, 9 (in two regions) and 10. In particular, QTLs for LossFF_T14_ on chr. 8 and 10 co-localized with those for FF_T14_. The QTL on chr. 9A had the strongest effect (LOD score = 22). For color components, QTLs were detected on chr. 5, 8, 9 (A and B), 11 (for L*), chr. 5, 9A, 11 (for a*_T14_), and chr. 8 and 11 (for b*_T14_). Both parents provided positive alleles depending on the QTLs. The QTL on chr. 11 had the strongest effect for all three components (with a LOD>11). Several overlapping QTL intervals were identified (**Figure S7B**). QTLs for color parameters were frequently mapped together, and in regions carrying FW (chr. 5), or FF (chr. 8, 9A, 9B) QTLs.

Overall, CIM identified numerous QTLs controlling fruit quality and shelf-life traits in both populations, with several genomic regions shared among traits or populations and others being population-specific, highlighting the complex genetic architecture underlying tomato shelf life.

#### Targeted BSA-Seq for shelf life traits

To validate and refine the QTLs identified by CIM, targeted BSA-Seq analyses were performed using extreme segregant pools selected for shelf-life related traits in each population. Average trait values of ENEA segregant pools were respectively +23%/-21% and +96%/-64% higher/lower than the F2 population average for FF_T14_ and LossFW_T14_ (**Figure S8 A** and **B**). BSA-Seq analysis allowed the identification of twelve QTLs for FF_T14_ and seven for LossFW_T14_ in the ENEA bulks (**Table 2**, **Figure 3 and S9**). The most notable QTLs identified by BSA-Seq were those on chr. 5 – one for LossFW_T14_ (58 - 61 Mb) and one for FF_T14_ (52 -64 Mb) – which overlapped with those obtained by CIM for FF traits (58 - 61 Mb), identifying a hotspot controlling shelf life in ENEA BSA-Seq analysis. Additional overlapping regions between CIM- and BSA-Seq-derived QTLs for LossFW_T14_ were found on chr. 2 and chr. 3. From the INRAE bulks (**Figure S8 C** and **D**), BSA-Seq identified four QTLs for FF_T0_ and six for LossFF_T14_ (**Table 2**, **Figure 3**). For FF_T0_, the four regions corresponded to QTLs mapped by CIM within the whole F2 population but covering much smaller intervals (except for the QTL on chr. 9B) (**Figure 3**). For LossFF_T14_, six QTLs were identified and only one (on chr. 10) was common to CIM and BSA-Seq. A region on chr. 7 co-localized with a QTL for FF_T0_. New QTLs were detected on chr. 4, 11, and 12. Most notably, QTLs for FF from both populations overlapped in two regions at both extremities of chr. 9 and on chr. 12, a common region was detected for LossFF_T14_ (INRAE) and FF_T14_ (ENEA) (**Figure 3**). In total, BSA-Seq allowed the detection of more QTLs with smaller intervals than in QTL mapping, as 48% of the BSA-Seq QTLs had an interval lower than 2.5 Mb, instead of 26% of the QTLs detected by CIM. Overall, targeted BSA-Seq complemented CIM by refining several QTL intervals and identifying additional genomic regions associated with shelf-life traits.

### Search for candidate genes in shelf life-related QTLs

A total of 60 QTLs for FW- and FF-related traits were identified by combined CIM and BSA-Seq mapping in the two F2 populations (**Table 2**). After merging overlapping intervals that likely represent the same underlying QTL, 26 QTL regions were retained, including 16 associated with FF-related traits, five with FW-related traits, and five with both trait classes (**Figure 3**; **Table S7**). No QTL overlapped the major ripening regulator *nor*, while *rin* was included in only one ENEA FF_T14_-associated QTL (**Figure 3**). This suggests that the loci identified here are unlikely to act through strong suppression of ripening, but rather through more specific effects on processes such as cell-wall remodeling, water balance, hormone signaling, primary metabolism and stress responses.

Candidate gene search was focused on FW, LossFW, FF and LossFF QTLs with confidence intervals smaller than 2.5 Mb, identified either by CIM or BSA-Seq. This led to the selection, in the ENEA population, of five FF_T14_ QTLs on chr. 2, 5, 9 and 12, three LossFW_T14_ QTLs on chromosomes 2, 3 and 5, and one FW QTL on chromosome 11. In the INRAE population, candidate searches were carried out for two FW QTLs on chromosomes 6 and 7, two FF_T0_ QTLs on chr. 2 and 7, and five LossFF_T14_ QTLs on chr. 4, 7, 10 and 12 (**Table 2; Tables S8** and **S9**). Candidate genes were selected according to their functional annotation, fruit expression in the TEA database, and the presence of coding or regulatory polymorphisms between the parental lines. The main candidate regions and biological processes are summarized in **Table 3**.

**Table 3.**
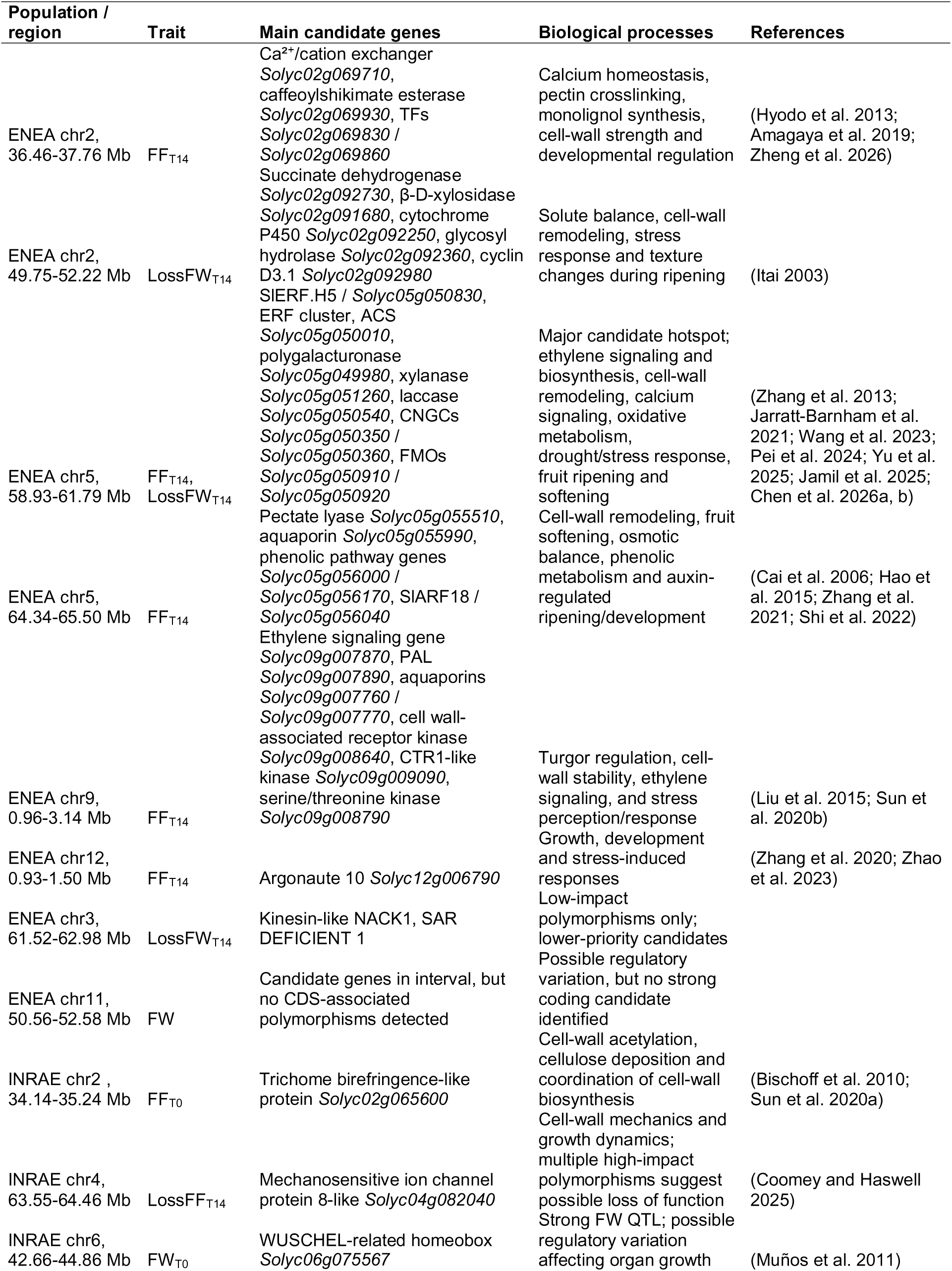
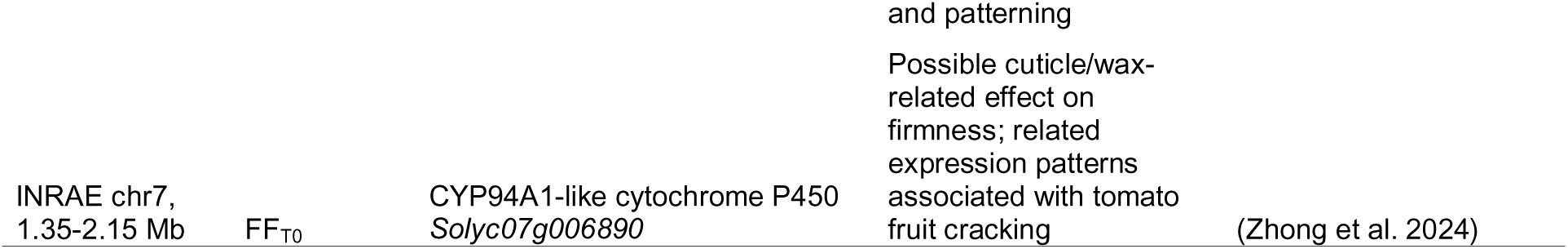
Main QTLs and candidate genes identified in the ENEA and INRAE populations for QTLs with confidence intervals lower than 2.5 Mb.

In the ENEA population, the most compelling candidate region was located on chr. 5, where overlapping FF and LossFW_T14_ QTLs defined a major interval between 58.9 and 61.8 Mb (**Table 3**). This region contained SlERF.H5 (*Solyc05g050830*), carrying one high-impact and two non-synonymous polymorphisms, together with additional ERF transcription factors, an ACC synthase, polygalacturonase, xylanase, laccase, cyclic nucleotide-gated channels and flavin-containing monooxygenases (**Table S8**). The co-localization of these QTLs with genes involved in ethylene signaling, cell-wall metabolism, calcium signaling and oxidative/stress responses supports this region as a major candidate genomic interval in the control of tomato fruit firmness and postharvest performance. A second ENEA chr. 5 interval contained additional candidates related to cell-wall remodeling and osmotic balance, including pectate lyase, aquaporin and SlARF18 genes (**Tables 3** and **S8**).

Additional ENEA candidate regions pointed to complementary mechanisms. On chr. 2, candidate genes included a Ca2+/cation exchanger, a caffeoylshikimate esterase, β-D-xylosidase, glycosyl hydrolase and cyclin D3.1, suggesting possible effects on calcium homeostasis, pectin crosslinking, cell-wall structure and texture changes during ripening. On chr. 9, candidates included genes related to ethylene signaling, aquaporin-mediated water transport, PAL activity, receptor kinase signaling and stress response, including a CTR1-like protein kinase. On chr. 12, a moderate-impact polymorphism was detected in Argonaute 10, a gene associated with developmental and stress-related regulation. By contrast, the chr. 3 LossFW_T14_ interval contained only low-impact polymorphisms, and no CDS-associated polymorphisms were detected in candidate genes within the chr. 11 FW interval (**Tables S8** and **3**).

In the INRAE population, three regions contained particularly compelling candidates (**Tables S9** and **3**). A high-impact polymorphism was detected in a trichome birefringence-like gene on chr. 2, potentially affecting cell-wall acetylation and cellulose deposition. On chr. 7, a strong FF_T0_ QTL contained a CYP94A1-like cytochrome P450 gene, possibly linked to cuticular wax metabolism and fruit surface properties. On chr. 4, the LossFF_T14_ QTL contained an MSL8-like mechanosensitive ion channel gene with several high-impact polymorphisms, suggesting a possible loss-of-function allele affecting cell-wall mechanics and growth dynamics. Finally, the strong chr. 6 FW QTL did not contain coding polymorphisms in the main candidate gene but may involve regulatory variation affecting a WUSCHEL-related homeobox gene, consistent with a role in organ growth and patterning. Overall, the candidate gene analysis suggests that natural variation in tomato shelf life-related traits is mediated by multiple partially independent biological processes. These include ethylene signaling, cell-wall remodeling, calcium homeostasis, water transport, cuticle-associated metabolism, mechanosensing and stress responses.

The combination of QTL mapping, BSA-Seq, parental resequencing and candidate prioritization therefore provides a focused set of loci for future functional validation and breeding applications (**Tables S7-S9; Table 3**).

### Screening for polymorphisms in QTLs common to the two populations on chr. 9 and 12

To further prioritize candidate genes, we searched for polymorphisms shared between the two parental pairs within the overlapping FF- and LossFF-associated QTLs identified on chromosomes 9 and 12. On chr. 9, a total of 1,378 and 1,497 SNPs were identified between the ENEA and INRAE parental lines, respectively, of which 295 were shared between the two populations. Among them, twenty-one SNPs in eleven genes had a non-synonymous impact (**Table S10**). The two most notable SNPs were in a CTR1-like protein kinase (*Solyc09g009090*) which acts as a negative regulator in the ethylene signaling pathway (Lin et al. 2008). Other polymorphisms were detected in *Solyc09g008750* (VQ motif-containing protein), *Solyc09g008913* (cytochrome P450) and *Solyc09g008790* (serine/threonine protein kinase) genes, which regulate diverse developmental processes, as well as responses to biotic and abiotic stresses. Plant P450 monoxygenases belong to a vast enzyme family with multiple roles in biosynthetic routes and development (Fang et al. 2024). Another cytochrome P450 gene, *SlKLUH*, was associated with the *fw3.2* locus controlling tomato fruit size, stressing the importance of this enzyme family in the control of economically-important agronomic traits in different species (Zhang et al. 2012; Chakrabarti et al. 2013).

The chr. 12 region was poorly polymorphic in ENEA population (only 66 SNPs between parental lines), in contrast to INRAE parents (5656 polymorphisms). Only 43 common polymorphisms were detected, among which one non-synonymous mutation in the *Argonaute 10* gene (*Solyc12g006790*) already discussed above (**Table S10**). The limited number of shared polymorphisms on chr. 12 markedly reduced the number of common candidate genes compared with chr. 9. Overall, comparison of the two populations reduced the number of high-priority candidate genes and highlighted a small set of shared coding polymorphisms that represent promising targets for future functional validation.

## Conclusion

This study shows that tomato shelf life should be treated as a dynamic postharvest trait rather than as a simple extension of fruit firmness at harvest. In two intraspecific F_2_ populations, fruit firmness at harvest was only weakly associated with firmness retention during storage, indicating that these traits should not be considered interchangeable proxies of tomato shelf life. This finding emphasizes that shelf life is a dynamic trait requiring time-resolved phenotyping, and highlighting the need for high-throughput phenotyping strategies adapted to postharvest fruit evolution. Our results further support the view that tomato shelf life is a complex trait resulting from the interaction of multiple biological processes, including cell-wall remodeling, hormone signaling, water balance, primary metabolism and stress responses.

Despite the use of intraspecific populations derived from *Solanum lycopersicum* crosses, characterized by extended regions of identity by descent and reduced polymorphism, we identified a substantial number of QTLs controlling fruit firmness and its postharvest evolution. While the total number of detected QTLs was lower than typically reported in interspecific studies, their detection is facilitated by simpler genetic backgrounds, making them particularly relevant for breeding applications. The limited overlap between QTLs for firmness at harvest and firmness loss during storage supports the hypothesis that these traits are, at least in part, under distinct genetic control and reinforces the polygenic nature of shelf life-related traits. Frequent co-localizations of QTLs for different traits within populations suggest either pleiotropic effects or tightly linked loci, consistent with previously-reported QTL clusters (Chapman et al. 2012).

Candidate gene analysis, while informative, remains constrained by current knowledge and the absence of transcriptomic data such as RNA-Seq from parental lines, which may have facilitated the identification of additional candidate regulators. From a methodological perspective, the comparison between CIM and BSA-Seq highlights the complementary value of sequencing-based approaches. BSA-Seq, benefiting from reduced sequencing costs, enabled the detection of additional QTLs and often narrowed confidence intervals, although it remains dependent on accurate and high-quality phenotyping.

Future work should prioritize the functional validation of promising candidate genes, for example through gene editing approaches, to confirm their role in the regulation of shelf life. In parallel, the integration of identified QTLs into marker-assisted selection or genomic prediction frameworks offers practical avenues for breeding tomato varieties with improved postharvest performance. Together, these results contribute to a better understanding of the complex genetic architecture of fruit shelf life and provide useful resources for future functional studies and breeding efforts aimed at improving postharvest performance in tomato.

## Supplementary Materials

### Supplementary Figures

**Suppl. Figure S1. Distribution charts of color space L*-related traits in studied F2 populations.** A-B: Distribution charts of L*_T0_ in the ENEA (A) and INRAE (B). C-D: Distribution charts of delta L*_T14_ in the ENEA (C) and INRAE (D). Blue histograms: F2 progeny bins. Dark blue dotted line: average of F2 population. Red line: average of P1. Green line: average of P2. Orange line: average of F1 hybrid. Black line: distribution curve of F2 population.

**Suppl. Figure S2. Distribution charts of color space a*-related traits in studied F2 populations.** A-B: Distribution charts of a*_T0_ in the ENEA (A) and INRAE (B). C-D: Distribution charts of delta a*_T14_ in the ENEA (C) and INRAE (D). Blue histograms: F2 progeny bins. Dark blue dotted line: average of F2 population. Red line: average of P1. Green line: average of P2. Orange line: average of F1 hybrid. Black line: distribution curve of F2 population.

**Suppl. Figure S3. Distribution charts of color space b*-related traits in studied F2 populations.** A-B: Distribution charts of b*_T0_ in the ENEA (A) and INRAE (B). C-D: Distribution charts of delta b*_T14_ in the ENEA (C) and INRAE (D). Blue histograms: F2 progeny bins. Dark blue dotted line: average of F2 population. Red line: average of P1. Green line: average of P2. Orange line: average of F1 hybrid. Black line: distribution curve of F2 population.

**Suppl. Figure S4. Relationships between shelf life-related traits in the two F2 populations.** Top panels: ENEA. Bottom panels: INRAE.

**Suppl. Figure S5. Polymorphism and SNP density distributions in parental lines of ENEA and INRAE populations.** A: Polymorphism distributions of Agriplex SNP panel in parental lines of ENEA (green) and INRAE (red) populations. Black lines: SNP marker position. B-C: SNP density distributions in genome sequences of parental lines of ENEA (B) and INRAE (C) populations.

**Suppl. Figure S6. Genetic maps constructed with polymorphic SNP markers.** dots indicate clusters of markers. A: ENEA map. B: INRAE map. Red dots indicate clusters of markers.

**Suppl. Figure S7. Genetic maps of QTLs detected in the ENEA and INRAE F2 populations by Composite Interval Mapping.** A: Map of the ENEA population. B: Map of the INRAE F2 population. QTLs for each trait (and experimental stage) are represented with bars of different colors. Bar length indicates the confidence interval of each QTL. FF: fruit firmness; LossFF_T14_: Loss of FF at T14; FW: fruit weight; LossFW_T7/T14_: Loss of FW at T7 or T14; L*: fruit skin brightness; a*: red/green color; b*: yellow/blue color.

**Suppl. Figure S8. Average trait values of selected F2 segregants used for BSA-Seq relative to the population distribution.** A-B: ENEA FF_T14_ (A) and LossFW_T14_ (B). C-D: INRAE FF_T0_ (C) and LossFF_T14_ (D). Blue histograms: F2 progeny bins. Dark blue dotted line: average of F2. Light green line: average of low-trait segregants. Dark green line: average of high-trait segregants.

**Suppl. Figure S9**. **BSA-Seq plots for extreme segregants of shelf life related traits in tomato.** (a) ENEA F_₂_ population for LossFW_T14_ (panel A) and FF_T14_ (panel B); (b) INRAE F_₂_ population for LossFF_T14_ (panel C) and FF_T0_ (panel D). Each panel shows the genome-wide distribution of the ΔSNP-index (blue dots) along the 12 chromosomes of *Solanum lycopersicum*, obtained by QTL-seq analysis (Sugihara et al., 2022; Takagi et al., 2013) of extreme bulk segregants. The red dashed lines indicate the position-specific (P<0.01) significance threshold estimated by simulation. Genomic regions where the ΔSNP-index exceeded the threshold were expanded based on linkage decay to a minimum r² of 0.2; Intervals narrower than 3.5 Mb were retained; overlapping intervals were merged and are shown as green shaded areas.

### Supplementary Tables

**Suppl. Table S1. Phenotypes of ENEA material for studied traits.** For each genotype, average trait values are reported (n=5 fruits). F2 segregants chosen for BSA-Seq analyses are identified in the Bulk column (1: low LossFW_T14_; 2: high LossFW_T14_; 3: low FF_T14_; 4: high FF_T14_).

**Suppl. Table S2. Phenotypes of INRAE material for studied traits.** For each genotype, average trait values are reported (n=5 fruits). F2 segregants chosen for BSA-Seq analyses are identified in the Bulk column (1: low FF_T0_; 2: high FF_T0_; 3: high LossFF_T14_; 4: low LossFF_T14_).

**Suppl. Table S3. Genotypes of ENEA materials used for genotyping by Agriplex 1078-SNP panel.** Genotypes of F2 progenies, P1, P2 and F1 plants are in columns. Reference (REF) and alternative (ALT) are indicated. Genotypes at all the AGRIPLEX SNPs (0: reference allele, 1: Alternative allele) are in rows.

**Suppl. Table S4. Genotypes of INRAE F2 plants based on genotyping by Agriplex 1078-SNP panel.** F2 progenies are in columns. Genotypes at all the AGRIPLEX SNPs (A: reference allele, B: Alternative allele) are in rows.

**Suppl. Table S5. QTL information of the genetic map of ENEA F2 population obtained by Composite Interval Mapping.** LOD threshold: 3.30. QTLs with C.I. <2.5 Mb further searched for candidate genes are highlighted in bold.

**Suppl. Table S6. QTL information of the genetic map of INRAE F2 population obtained by Composite Interval Mapping.** LOD threshold: 3.30. QTLs with C.I. <2.5 Mb further searched for candidate genes are highlighted in bold.

**Suppl. Table S7. Characteristics of the 26 regions with overlapping QTLs of shelf life-related traits in studied populations.**

**Suppl. Table S8. Candidate genes for shelf life-related traits from ENEA genetic map and BSA-Seq QTLs with confidence interval <2.5Mb.** Expression data (in rpkm) were retrieved from https://tea.solgenomics.net/. MG: mature green. T.F.: Transcription factor. Polymorphisms were classified according to their impact on the protein sequence: HIGH for frameshift deletion and insertion, and stop gain; MODERATE for non-frameshift deletion or insertion, and non-synonymous modification; LOW for synonymous variation; MODIFIER for polymorphism in non-coding sequence.

**Suppl. Table S9. Candidate genes for shelf life-related QTLs from INRAE genetic map and BSA-Seq QTLs with confidence interval <2.5Mb.** Expression data (in rpkm) were retrieved from https://tea.solgenomics.net/ doi:10.193/bioinformatics/btx190. MG: mature green. Polymorphisms were classified according to their impact on the protein sequence: HIGH for frameshift deletion and insertion, and stop gain; MODERATE for non frameshift deletion or insertion, and non-synonymous modification; LOW for synonymous variation; MODIFIER for polymorphism in non-coding sequence.

**Suppl. Table S10. Common SNPs in INRAE and ENEA populations with moderate impact in two overlapping QTLs regions on chromosomes 9 and 12.**

## Acknowledgments

Authors thank Yolande Carretero and Mila Demaria for technical assistance, Tiziana Maria Sirangelo for contribution in early phases of the project and the experimental Unit A2M for greenhouse management. The experiments in this study comply with the current EU laws and ethics requirements by Projects’ Consortia.

## Statements and Declarations

### Data Availability

The genotype and phenotype datasets generated during and/or analyzed during the current study are available in supplementary tables. The sequences of BSA-Seq can be downloaded from BioProject ID: PRJNA1475219

### Authors’ contribution

GG, MC conceived experimental design; CR, GG, MC, wrote the manuscript; CR, CS, FT, KP, MB, PF, RD carried out experimental work; CR, FT, GA, MC, FB performed data analyses; all authors revised and approved the submitted manuscript.

## References

Albert E, Segura V, Gricourt J, et al (2016) Association mapping reveals the genetic architecture of tomato response to water deficit: focus on major fruit quality traits. J Exp Bot 67:6413–6430. 10.1093/jxb/erw411

Alpert KB, Grandillo S, Tanksley SD (1995) fw 2.2:a major QTL controlling fruit weight is common to both red- and green-fruited tomato species. Theor Appl Genet 91:994– 1000. 10.1007/BF00223911

Amagaya K, Shibuya T, Nishiyama M, et al (2019) Characterization and Expression Analysis of the Ca2+/Cation Antiporter Gene Family in Tomatoes. Plants 9:25. 10.3390/plants9010025

Arah IK, Amaglo H, Kumah EK, Ofori H (2015) Preharvest and Postharvest Factors Affecting the Quality and Shelf Life of Harvested Tomatoes: A Mini Review. Int J Agron 2015:1–6. 10.1155/2015/478041

Ballester A-R, Molthoff J, De Vos R, et al (2009) Biochemical and Molecular Analysis of Pink Tomatoes: Deregulated Expression of the Gene Encoding Transcription Factor SlMYB12 Leads to Pink Tomato Fruit Color. Plant Physiol 152:71–84. 10.1104/pp.109.147322

Bassolino L, Zhang Y, Schoonbeek H, et al (2013) Accumulation of anthocyanins in tomato skin extends shelf life. New Phytol 200:650–655. 10.1111/nph.12524

Bischoff V, Nita S, Neumetzler L, et al (2010) *TRICHOME BIREFRINGENCE* and Its Homolog *AT5G01360* Encode Plant-Specific DUF231 Proteins Required for Cellulose Biosynthesis in Arabidopsis. Plant Physiol 153:590–602. 10.1104/pp.110.153320

Broman KW, Wu H, Sen Ś, Churchill GA (2003) R/qtl: QTL mapping in experimental crosses. Bioinformatics 19:889–890. 10.1093/bioinformatics/btg112

Cai C, Xu C, Li X, et al (2006) Accumulation of lignin in relation to change in activities of lignification enzymes in loquat fruit flesh after harvest. Postharvest Biol Technol 40:163–169. 10.1016/j.postharvbio.2005.12.009

Cambiaso V, Gimenez MD, Da Costa JHP, et al (2019) Selected genome regions for fruit weight and shelf life in tomato RILs discernible by markers based on genomic sequence information. Breed Sci 69:447–454. 10.1270/jsbbs.19015

Carrari F (2006) Metabolic regulation underlying tomato fruit development. J Exp Bot 57:1883–1897. 10.1093/jxb/erj020

Causse M (2002) QTL analysis of fruit quality in fresh market tomato: a few chromosome regions control the variation of sensory and instrumental traits. J Exp Bot 53:2089– 2098. 10.1093/jxb/erf058

Chakrabarti M, Zhang N, Sauvage C, et al (2013) A cytochrome P450 regulates a domestication trait in cultivated tomato. Proc Natl Acad Sci 110:17125–17130. 10.1073/pnas.1307313110

Chapman NH, Bonnet J, Grivet L, et al (2012) High-Resolution Mapping of a Fruit Firmness-Related Quantitative Trait Locus in Tomato Reveals Epistatic Interactions Associated with a Complex Combinatorial Locus. Plant Physiol 159:1644–1657. 10.1104/pp.112.200634

Chen S (2023) Ultrafast one-pass FASTQ data preprocessing, quality control, and deduplication using fastp. iMeta 2:e107. 10.1002/imt2.107

Chen Y, Liao X, Wang X, et al (2026a) Genome-wide identification, classification and expression analysis of laccase family members in Solanum lycopersicum. BMC Plant Biol 26:337. 10.1186/s12870-026-08166-w

Chen Z, Liu J, Zhimo VY, et al (2026b) Biochemical and molecular regulation of tomato ripening and disease defense: A trade-off between quality and postharvest integrity. Food Chem 508:148476. 10.1016/j.foodchem.2026.148476

Çolak NG, Eken NT, Ülger M, et al (2023) Mapping of quantitative trait loci for the nutritional value of fresh market tomato. Funct Integr Genomics 23:121. 10.1007/s10142-023-01045-9

Coomey JH, Haswell ES (2025) Mechanosensitive ion channel MSL8 is required for oscillatory growth and cell wall dynamics in Arabidopsis pollen tubes. Plant Reprod 38:22. 10.1007/s00497-025-00530-4

Del Giudice R, Raiola A, Tenore GC, et al (2015) Antioxidant bioactive compounds in tomato fruits at different ripening stages and their effects on normal and cancer cells. J Funct Foods 18:83–94. 10.1016/j.jff.2015.06.060

Desaint H, Héreil A, Belinchon-Moreno J, et al (2024) Integration of QTL and transcriptome approaches for the identification of genes involved in tomato response to nitrogen deficiency. J Exp Bot 75:5880–5896. 10.1093/jxb/erae265

Diouf IA, Derivot L, Bitton F, et al (2018) Water Deficit and Salinity Stress Reveal Many Specific QTL for Plant Growth and Fruit Quality Traits in Tomato. Front Plant Sci 9:279. 10.3389/fpls.2018.00279

Diretto G, Frusciante S, Fabbri C, et al (2020) Manipulation of β_-_carotene levels in tomato fruits results in increased ABA content and extended shelf life. Plant Biotechnol J 18:1185–1199. 10.1111/pbi.13283

Fang Y, Tai Z, Hu K, et al (2024) Comprehensive Review on Plant Cytochrome P450 Evolution: Copy Number, Diversity, and Motif Analysis From Chlorophyta to Dicotyledoneae. Genome Biol Evol 16:evae240. 10.1093/gbe/evae240

Fernández-Muñoz R, Heredia A, Domínguez E (2022) The role of cuticle in fruit shelf-life. Curr Opin Biotechnol 78:102802. 10.1016/j.copbio.2022.102802

Fernandez-Pozo N, Zheng Y, Snyder SI, et al (2017) The Tomato Expression Atlas. Bioinformatics 33:2397–2398. 10.1093/bioinformatics/btx190

Frary A, Nesbitt TC, Frary A, et al (2000) fw2.2: A Quantitative Trait Locus Key to the Evolution of Tomato Fruit Size. Science 289:85–88. 10.1126/science.289.5476.85

Fumelli L, Ilyas M, Olivieri F, et al (2026) Knockout of a pectate lyase in tomato increases fruit firmness and reduces foliar susceptibility to pathogens. Plant Sci 363:112868. 10.1016/j.plantsci.2025.112868

Gao L, Gonda I, Sun H, et al (2019) The tomato pan-genome uncovers new genes and a rare allele regulating fruit flavor. Nat Genet 51:1044–1051. 10.1038/s41588-019-0410-2

Hao Y, Hu G, Breitel D, et al (2015) Auxin Response Factor SlARF2 Is an Essential Component of the Regulatory Mechanism Controlling Fruit Ripening in Tomato. PLOS Genet 11:e1005649. 10.1371/journal.pgen.1005649

Huang Z, Van Der Knaap E (2011) Tomato fruit weight 11.3 maps close to fasciated on the bottom of chromosome 11. Theor Appl Genet 123:465–474. 10.1007/s00122-011-1599-3

Hyodo H, Terao A, Furukawa J, et al (2013) Tissue Specific Localization of Pectin–Ca2+ Cross-Linkages and Pectin Methyl-Esterification during Fruit Ripening in Tomato (Solanum lycopersicum). PLoS ONE 8:e78949. 10.1371/journal.pone.0078949

Itai A (2003) Characterization of expression, and cloning, of -D-xylosidase and -L-arabinofuranosidase in developing and ripening tomato (Lycopersicon esculentum Mill.) fruit. J Exp Bot 54:2615–2622. 10.1093/jxb/erg291

Jamil A, Habib MA, Ibrahim ABM, et al (2025) A genome-wide approach to the comprehensive analysis of the serine carboxypeptidase-like protein family in Solanum lysopersicum L. Genet Resour Crop Evol 72:10185–10211. 10.1007/s10722-025-02559-w

Jarratt-Barnham E, Wang L, Ning Y, Davies JM (2021) The Complex Story of Plant Cyclic Nucleotide-Gated Channels. Int J Mol Sci 22:874. 10.3390/ijms22020874

Ji D, Liu W, Jiang L, Chen T (2023) Cuticles and postharvest life of tomato fruit: A rigid cover for aerial epidermis or a multifaceted guard of freshness? Food Chem 411:135484. 10.1016/j.foodchem.2023.135484

Juan-Cabot A, Galmés J, Conesa MÀ (2022) The tomato long shelf-life fruit phenotype: Knowledge, uncertainties and prospects. Sci Hortic 291:110578. 10.1016/j.scienta.2021.110578

Kapoor L, Simkin AJ, George Priya Doss C, Siva R (2022) Fruit ripening: dynamics and integrated analysis of carotenoids and anthocyanins. BMC Plant Biol 22:27. 10.1186/s12870-021-03411-w

Kaur G, Abugu M, Tieman D (2023) The dissection of tomato flavor: biochemistry, genetics, and omics. Front Plant Sci 14:1144113. 10.3389/fpls.2023.1144113

Kayikci HC, Aydin S, Adak A, et al (2025) Association mapping of tomato fruit quality for weight, firmness, brix, and color using GWAS. BMC Plant Biol 26:41. 10.1186/s12870-025-07838-3

Kim M, Nguyen TTP, Ahn J-H, et al (2021) Genome-wide association study identifies QTL for eight fruit traits in cultivated tomato ( *Solanum lycopersicum* L.). Hortic Res 8:203. 10.1038/s41438-021-00638-4

Lecomte L, Saliba-Colombani V, Gautier A, et al (2004) Fine mapping of QTLs of chromosome 2 affecting the fruit architecture and composition of tomato. Mol Breed 13:1–14. 10.1023/B:MOLB.0000012325.77844.0c

Li R, Sun S, Wang H, et al (2020) FIS1 encodes a GA2-oxidase that regulates fruit firmness in tomato. Nat Commun 11:5844. 10.1038/s41467-020-19705-w

Lin Z, Alexander L, Hackett R, Grierson D (2008) LeCTR2, a CTR1_-_like protein kinase from tomato, plays a role in ethylene signalling, development and defence. Plant J 54:1083–1093. 10.1111/j.1365-313X.2008.03481.x

Liu J, Van Eck J, Cong B, Tanksley SD (2002) A new class of regulatory genes underlying the cause of pear-shaped tomato fruit. Proc Natl Acad Sci 99:13302–13306. 10.1073/pnas.162485999

Liu M, Pirrello J, Chervin C, et al (2015) Ethylene control of fruit ripening: revisiting the complex network of transcriptional regulation. Plant Physiol pp.01361.2015. 10.1104/pp.15.01361

Majeed A, Johar P, Raina A, et al (2022) Harnessing the potential of bulk segregant analysis sequencing and its related approaches in crop breeding. Front Genet 13:944501. 10.3389/fgene.2022.944501

Mubarok S, Qonit MAH, Rahmat BPN, et al (2023) An overview of ethylene insensitive tomato mutants: Advantages and disadvantages for postharvest fruit shelf-life and future perspective. Front Plant Sci 14:1079052. 10.3389/fpls.2023.1079052

Muños S, Ranc N, Botton E, et al (2011) Increase in Tomato Locule Number Is Controlled by Two Single-Nucleotide Polymorphisms Located Near *WUSCHEL*. Plant Physiol 156:2244–2254. 10.1104/pp.111.173997

Ning Y, Wei K, Li S, et al (2023) Fine Mapping of fw6.3, a Major-Effect Quantitative Trait Locus That Controls Fruit Weight in Tomato. Plants 12:. 10.3390/plants12112065

Oeller PW, Lu M-W, Taylor LP, et al (1991) Reversible Inhibition of Tomato Fruit Senescence by Antisense RNA. Science 254:437–439. 10.1126/science.1925603

Oladipo EK, Oladipo B, Ojumu TV, Caleb OJ (2026) Postharvest losses of tomato: Causes, spoilage mechanisms, and advances in conventional and emerging technologies for preservation. Food Res Int 227:118247. 10.1016/j.foodres.2025.118247

Ortega_-_Salazar I, Crum D, Sbodio AO, et al (2024) Double CRISPR knockout of pectin degrading enzymes improves tomato shelf_-_life while ensuring fruit quality. PLANTS PEOPLE PLANET 6:330–340. 10.1002/ppp3.10445

Pascual L, Desplat N, Huang BE, et al (2015) Potential of a tomato MAGIC population to decipher the genetic control of quantitative traits and detect causal variants in the resequencing era. Plant Biotechnol J 13:565–577. 10.1111/pbi.12282

Pei Y, Xue Q, Shu P, et al (2024) Bifunctional transcription factors SlERF.H5 and H7 activate cell wall and repress gibberellin biosynthesis genes in tomato via a conserved motif. Dev Cell 59:1345–1359.e6. 10.1016/j.devcel.2024.03.006

Ranc N, Muños S, Xu J, et al (2012) Genome-Wide Association Mapping in Tomato (*Solanum lycopersicum*) Is Possible Using Genome Admixture of *Solanum lycopersicum* var. *cerasiforme*. G3 GenesGenomesGenetics 2:853–864. 10.1534/g3.112.002667

Rodríguez GR, Muños S, Anderson C, et al (2011) Distribution of *SUN, OVATE, LC*, and *FAS* in the Tomato Germplasm and the Relationship to Fruit Shape Diversity. Plant Physiol 156:275–285. 10.1104/pp.110.167577

Saliba-Colombani V, Causse M, Langlois D, et al (2001) Genetic analysis of organoleptic quality in fresh market tomato. 1. Mapping QTLs for physical and chemical traits: Theor Appl Genet 102:259–272. 10.1007/s001220051643

Shi Y, Vrebalov J, Zheng H, et al (2021) A tomato LATERAL ORGAN BOUNDARIES transcription factor, *SlLOB1*, predominantly regulates cell wall and softening components of ripening. Proc Natl Acad Sci 118:e2102486118. 10.1073/pnas.2102486118

Shi L, Liu Q, Qiao Q, et al (2022) Exploring the effects of pectate and pectate lyase on the fruit softening and transcription profiling of Solanum lycopersicum. Food Control 133:108636. 10.1016/j.foodcont.2021.108636

Sim S-C, Durstewitz G, Plieske J, et al (2012) Development of a Large SNP Genotyping Array and Generation of High-Density Genetic Maps in Tomato. PLoS ONE 7:e40563. 10.1371/journal.pone.0040563

Sugihara Y, Young L, Yaegashi H, et al (2022) High-performance pipeline for MutMap and QTL-seq. PeerJ 10:e13170. 10.7717/peerj.13170

Sun A, Yu B, Zhang Q, et al (2020a) MYC2-Activated *TRICHOME BIREFRINGENCE-LIKE37* Acetylates Cell Walls and Enhances Herbivore Resistance. Plant Physiol 184:1083–1096. 10.1104/pp.20.00683

Sun Z, Song Y, Chen D, et al (2020b) Genome-Wide Identification, Classification, Characterization, and Expression Analysis of the Wall-Associated Kinase Family during Fruit Development and under Wound Stress in Tomato (Solanum lycopersicum L.). Genes 11:1186. 10.3390/genes11101186

Takagi H, Akira Abe A, Yoshida K, et al. (2013) QTL _-_seq: Rapid Mapping of Quantitative Trait Loci in Rice by Whole Genome Resequencing of DNA from Two Bulked Populations. Plant J 74:174–83. 10.1111/tpj.12105

The Tomato Genome Consortium (2012) The tomato genome sequence provides insights into fleshy fruit evolution. Nature 485:635–641. 10.1038/nature11119

Uluisik S, Oney-Birol S (2021) Uncovering candidate genes involved in postharvest ripening of tomato using the Solanum pennellii introgression line population by integrating phenotypic data, RNA-Seq, and SNP analyses. Sci Hortic 288:110321. 10.1016/j.scienta.2021.110321

Vicente AR, Saladié M, Rose JK, Labavitch JM (2007) The linkage between cell wall metabolism and fruit softening: looking to the future. J Sci Food Agric 87:1435– 1448. 10.1002/jsfa.2837

Wang K, Li M, Hakonarson H (2010) ANNOVAR: functional annotation of genetic variants from high-throughput sequencing data. Nucleic Acids Res 38:e164–e164. 10.1093/nar/gkq603

Wang R, Lammers M, Tikunov Y, et al (2020) The rin, nor and Cnr spontaneous mutations inhibit tomato fruit ripening in additive and epistatic manners. Plant Sci 294:110436. 10.1016/j.plantsci.2020.110436

Wang L, Zhou Y, Ding Y, et al (2023) Novel flavin-containing monooxygenase protein FMO1 interacts with CAT2 to negatively regulate drought tolerance through ROS homeostasis and ABA signaling pathway in tomato. Hortic Res 10:uhad037. 10.1093/hr/uhad037

Yogendra KN, Ramanjini Gowda PH (2013) Phenotypic and molecular characterization of a tomato (Solanum lycopersicum L.) F2 population segregation for improving shelf life. Genet Mol Res 12:506–518. 10.4238/2013.January.9.4

Yu S, Wang H, Garcia-Caparros P, Liu M (2025) Revisiting the functions of ethylene response factors (ERFs) in tomato. Plant Horm 1:0–0. 10.48130/ph-0025-0008

Zhang N, Brewer MT, Van Der Knaap E (2012) Fine mapping of fw3.2 controlling fruit weight in tomato. Theor Appl Genet 125:273–284. 10.1007/s00122-012-1832-8

Zhang T, Wang X, Lu Y, et al (2013) Genome-Wide Analysis of the Cyclin Gene Family in Tomato. Int J Mol Sci 15:120–140. 10.3390/ijms15010120

Zhang L, Zhu M, Ren L, et al (2018) The SlFSR gene controls fruit shelf-life in tomato. J Exp Bot 69:2897–2909. 10.1093/jxb/ery116

Zhang C, Fan L, Le BH, et al (2020) Regulation of ARGONAUTE10 Expression Enables Temporal and Spatial Precision in Axillary Meristem Initiation in Arabidopsis. Dev Cell 55:603–616.e5. 10.1016/j.devcel.2020.10.019

Zhang M, Yang H, Zhu F, et al (2021) Transcript profiles analysis of citrus aquaporins in response to fruit water loss during storage. Plant Biol 23:819–830. 10.1111/plb.13269

Zhang S, Wu S, Jia Z, et al (2024) Exploring the influence of a single_-_nucleotide mutation in EIN4 on tomato fruit firmness diversity through fruit pericarp microstructure. Plant Biotechnol J 22:2379–2394. 10.1111/pbi.14352

Zhao J, Sauvage C, Zhao J, et al (2019) Meta-analysis of genome-wide association studies provides insights into genetic control of tomato flavor. Nat Commun 10:1534. 10.1038/s41467-019-09462-w

Zhao Z, Yang S-J, Yin X-X, et al (2023) ARGONAUTE 1: a node coordinating plant disease resistance with growth and development. Phytopathol Res 5:38. 10.1186/s42483-023-00194-w

Zheng X, Zhang X, Liu W, et al (2026) Lignin in fruits: Structure, function, and frontier perspectives on biosynthesis and regulation. Food Res Int 225:117943. 10.1016/j.foodres.2025.117943

Zhong Z, Wu Z, Zhou R, et al (2024) Ribo-seq and RNA-seq analyses enrich the regulatory network of tomato fruit cracking. BMC Plant Biol 24:1214. 10.1186/s12870-024-05937-1

Zhou Y, Li Z, Su X, et al (2024) Histone deacetylase SlHDA7 impacts fruit ripening and shelf life in tomato. Hortic Res 11:uhae234. 10.1093/hr/uhae234

