## Supplementary material for "Postharvest tomato shelf life is genetically distinct from fruit firmness: evidence from two F_2_ populations": supp figs

### Slide 1
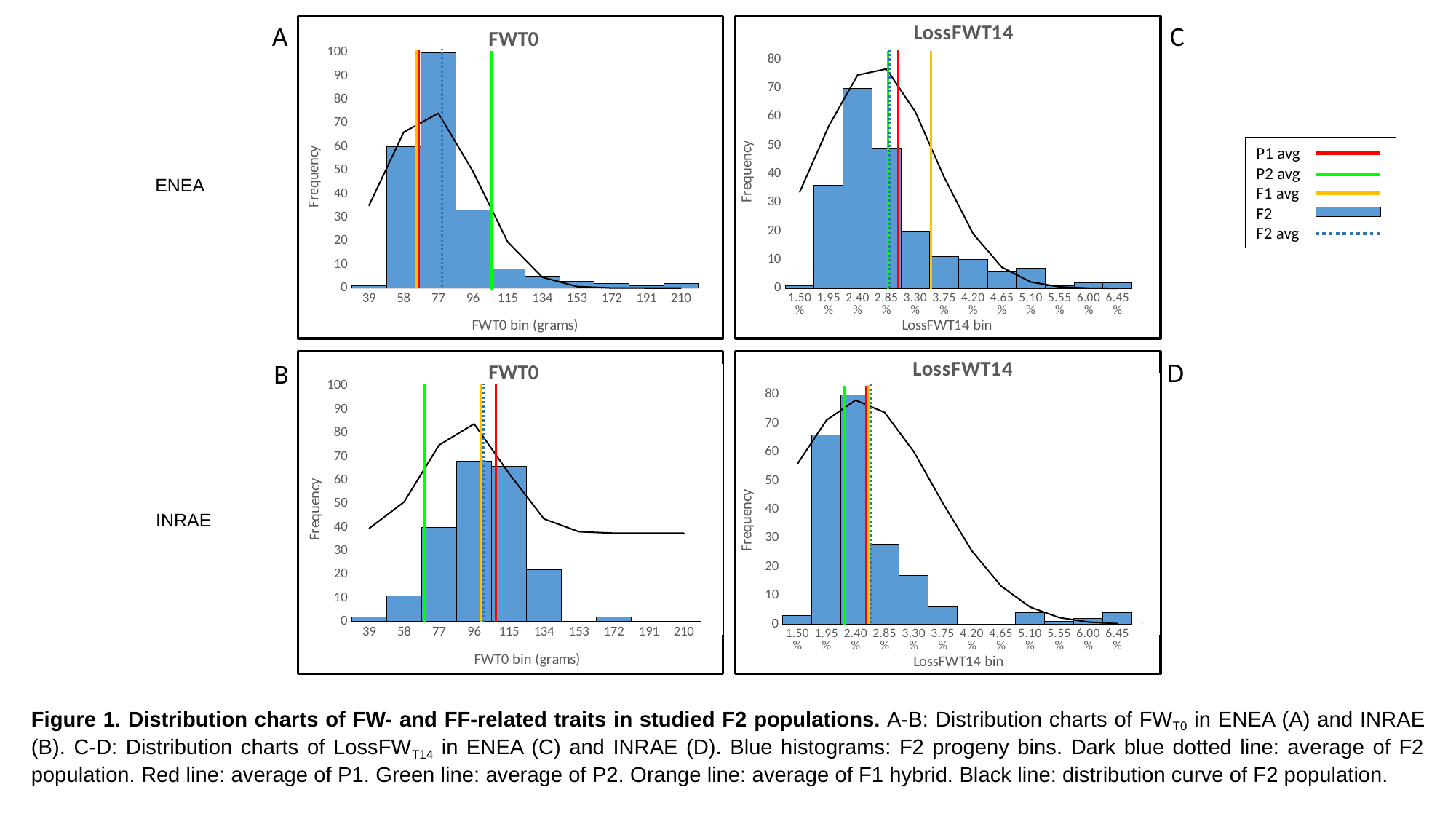

C
A
###### Chart: FWT0
| Category | | |
|---|---|---|
| 39 | 1.0 | 0.0070136445411499585 |
| 58 | 60.0 | 0.013232379053837768 |
| 77 | 100.0 | 0.014839116319517901 |
| 96 | 33.0 | 0.009891315840662138 |
| 115 | 8.0 | 0.003919006867025641 |
| 134 | 5.0 | 0.0009229409248011966 |
| 153 | 3.0 | 0.00012919559534507922 |
| 172 | 2.0 | 1.0749727017214246e-05 |
| 191 | 1.0 | 5.316466582457922e-07 |
| 210 | 2.0 | 1.5628765160770488e-08 |
###### Chart: LossFWT14
| Category | | |
|---|---|---|
| 1.4999999999999999E-2 | 1.0 | 18.968642661259963 |
| 1.95E-2 | 36.0 | 31.893124720436816 |
| 2.4E-2 | 70.0 | 41.94418126384187 |
| 2.8500000000000001E-2 | 49.0 | 43.147953910733 |
| 3.3000000000000002E-2 | 20.0 | 34.71862564694778 |
| 3.7499999999999999E-2 | 11.0 | 21.85137037822757 |
| 4.2000000000000003E-2 | 10.0 | 10.757433396593816 |
| 4.65E-2 | 6.0 | 4.142404336458472 |
| 5.0999999999999997E-2 | 7.0 | 1.2476999103746926 |
| 5.5500000000000001E-2 | 1.0 | 0.29395552975771005 |
| 0.06 | 2.0 | 0.05417101326642746 |
| 6.4500000000000002E-2 | 2.0 | 0.007808473094536076 |
P1 avg
P2 avg
F1 avg
F2
F2 avg
ENEA
###### Chart: FWT0
| Category | | |
|---|---|---|
| 39 | 2.0 | 0.0008358793755489122 |
| 58 | 11.0 | 0.0053318515722205325 |
| 77 | 40.0 | 0.014988368166687714 |
| 96 | 68.0 | 0.018568315788596218 |
| 115 | 66.0 | 0.010137538918391349 |
| 134 | 22.0 | 0.0024391270094665076 |
| 153 | 0.0 | 0.00025862955708143824 |
| 172 | 2.0 | 1.2085475898805591e-05 |
| 191 | 0.0 | 2.4888069853034504e-07 |
| 210 | 0.0 | 2.258710325978236e-09 |
###### Chart: LossFWT14
| Category | | |
|---|---|---|
| 1.4999999999999999E-2 | 3.0 | 24.44630919768964 |
| 1.95E-2 | 66.0 | 31.13516988711643 |
| 2.4E-2 | 80.0 | 34.16863978021521 |
| 2.8500000000000001E-2 | 28.0 | 32.31042361990571 |
| 3.3000000000000002E-2 | 17.0 | 26.326681842398695 |
| 3.7499999999999999E-2 | 6.0 | 18.483665871691304 |
| 4.2000000000000003E-2 | 0.0 | 11.181977256151203 |
| 4.65E-2 | 0.0 | 5.828913508780369 |
| 5.0999999999999997E-2 | 4.0 | 2.618153642960971 |
| 5.5500000000000001E-2 | 1.0 | 1.0133072687478497 |
| 0.06 | 2.0 | 0.3379291828931423 |
| 6.4500000000000002E-2 | 4.0 | 0.0971066002743281 |D
B
INRAE
Figure 1. Distribution charts of FW- and FF-related traits in studied F2 populations. A-B: Distribution charts of FW_T0_ in ENEA (A) and INRAE (B). C-D: Distribution charts of LossFW_T14_ in ENEA (C) and INRAE (D). Blue histograms: F2 progeny bins. Dark blue dotted line: average of F2 population. Red line: average of P1. Green line: average of P2. Orange line: average of F1 hybrid. Black line: distribution curve of F2 population.

### Slide 2
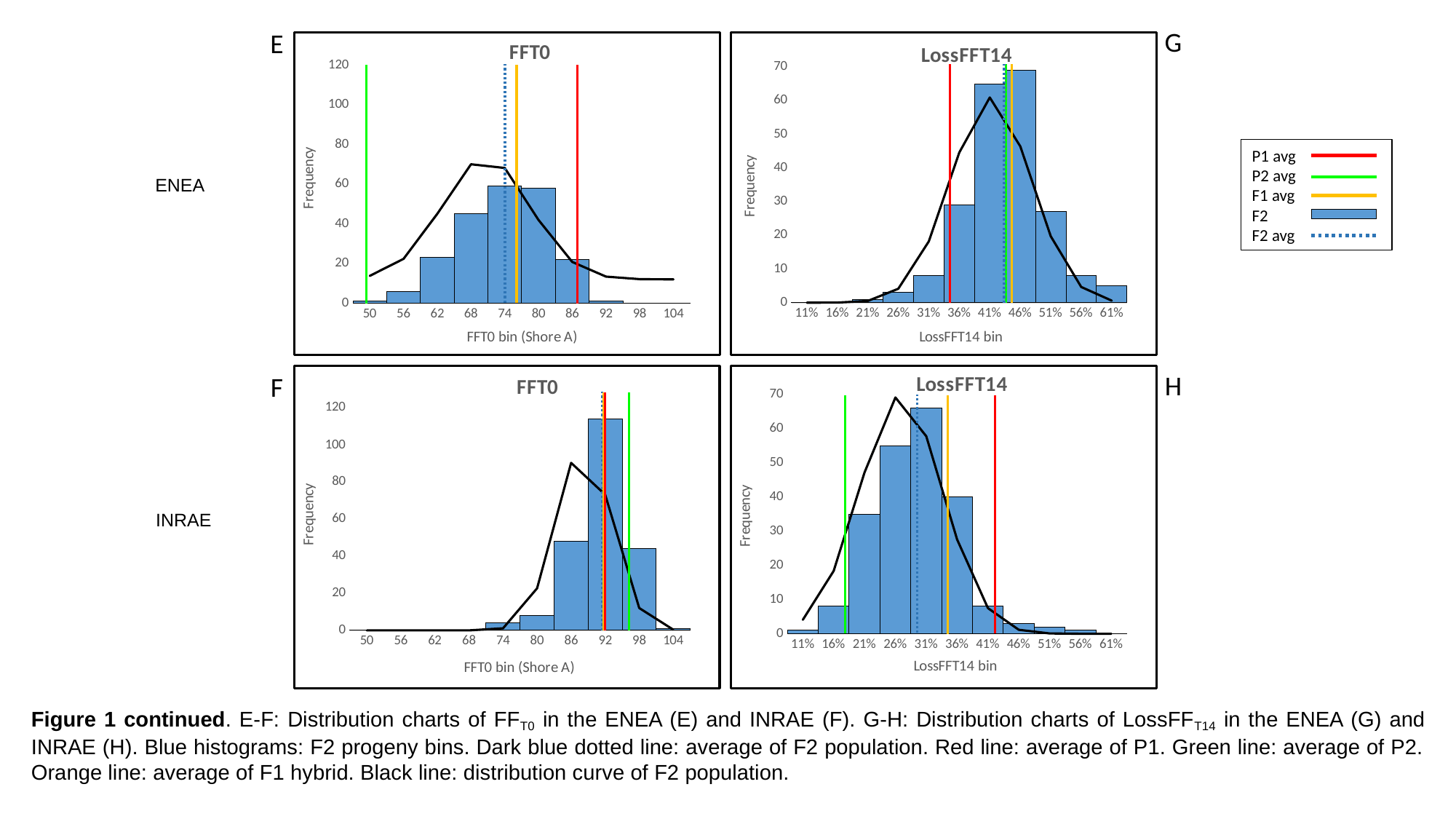

G
E
###### Chart: FFT0
| Category | | |
|---|---|---|
| 50 | 1.0 | 0.0014941404034773996 |
| 56 | 6.0 | 0.008641473683996722 |
| 62 | 23.0 | 0.027542920694826247 |
| 68 | 45.0 | 0.04837911834096214 |
| 74 | 59.0 | 0.04683081171200221 |
| 80 | 58.0 | 0.02498222982082083 |
| 86 | 22.0 | 0.007344402861186905 |
| 92 | 1.0 | 0.0011898920446819152 |
| 98 | 0.0 | 0.00010623910631853154 |
| 104 | 0.0 | 5.227415659872236e-06 |
###### Chart: LossFFT14
| Category | | |
|---|---|---|
| 0.11 | 0.0 | 0.00014735689868369863 |
| 0.16 | 0.0 | 0.0037307044940022823 |
| 0.21 | 1.0 | 0.052676440242474185 |
| 0.26 | 3.0 | 0.4148080515838925 |
| 0.31 | 8.0 | 1.8217262961710319 |
| 0.36 | 29.0 | 4.4619455651512245 |
| 0.41 | 65.0 | 6.0949563363398775 |
| 0.46 | 69.0 | 4.643250567183824 |
| 0.51 | 27.0 | 1.9727808015792503 |
| 0.56000000000000005 | 8.0 | 0.4674559972599548 |
| 0.61 | 5.0 | 0.061774299556264484 |
P1 avg
P2 avg
F1 avg
F2
F2 avg
ENEA
H
F
###### Chart: FFT0
| Category | | |
|---|---|---|
| 50 | 0.0 | 8.099551607089774e-16 |
| 56 | 0.0 | 9.219892794379798e-12 |
| 62 | 0.0 | 2.136565981055834e-08 |
| 68 | 0.0 | 1.0079344534756447e-05 |
| 74 | 4.0 | 0.0009679965637530389 |
| 80 | 8.0 | 0.018925217709712245 |
| 86 | 48.0 | 0.07532402108996836 |
| 92 | 114.0 | 0.061031163853857255 |
| 98 | 44.0 | 0.010066890649691781 |
| 104 | 1.0 | 0.00033803727642988963 |
###### Chart: LossFFT14
| Category | | |
|---|---|---|
| 0.11 | 1.0 | 0.350106009256822 |
| 0.16 | 8.0 | 1.5746548162527152 |
| 0.21 | 35.0 | 4.041890717227551 |
| 0.26 | 55.0 | 5.9210287358057805 |
| 0.31 | 66.0 | 4.950204846044718 |
| 0.36 | 40.0 | 2.3619058055715114 |
| 0.41 | 8.0 | 0.6431545950355917 |
| 0.46 | 3.0 | 0.09994971968447967 |
| 0.51 | 2.0 | 0.008864641659657556 |
| 0.56000000000000005 | 1.0 | 0.00044869805726020215 |
| 0.61 | 0.0 | 1.2961659076013753e-05 |
INRAE
Figure 1 continued. E-F: Distribution charts of FF_T0_ in the ENEA (E) and INRAE (F). G-H: Distribution charts of LossFF_T14_ in the ENEA (G) and INRAE (H). Blue histograms: F2 progeny bins. Dark blue dotted line: average of F2 population. Red line: average of P1. Green line: average of P2. Orange line: average of F1 hybrid. Black line: distribution curve of F2 population.

### Slide 3
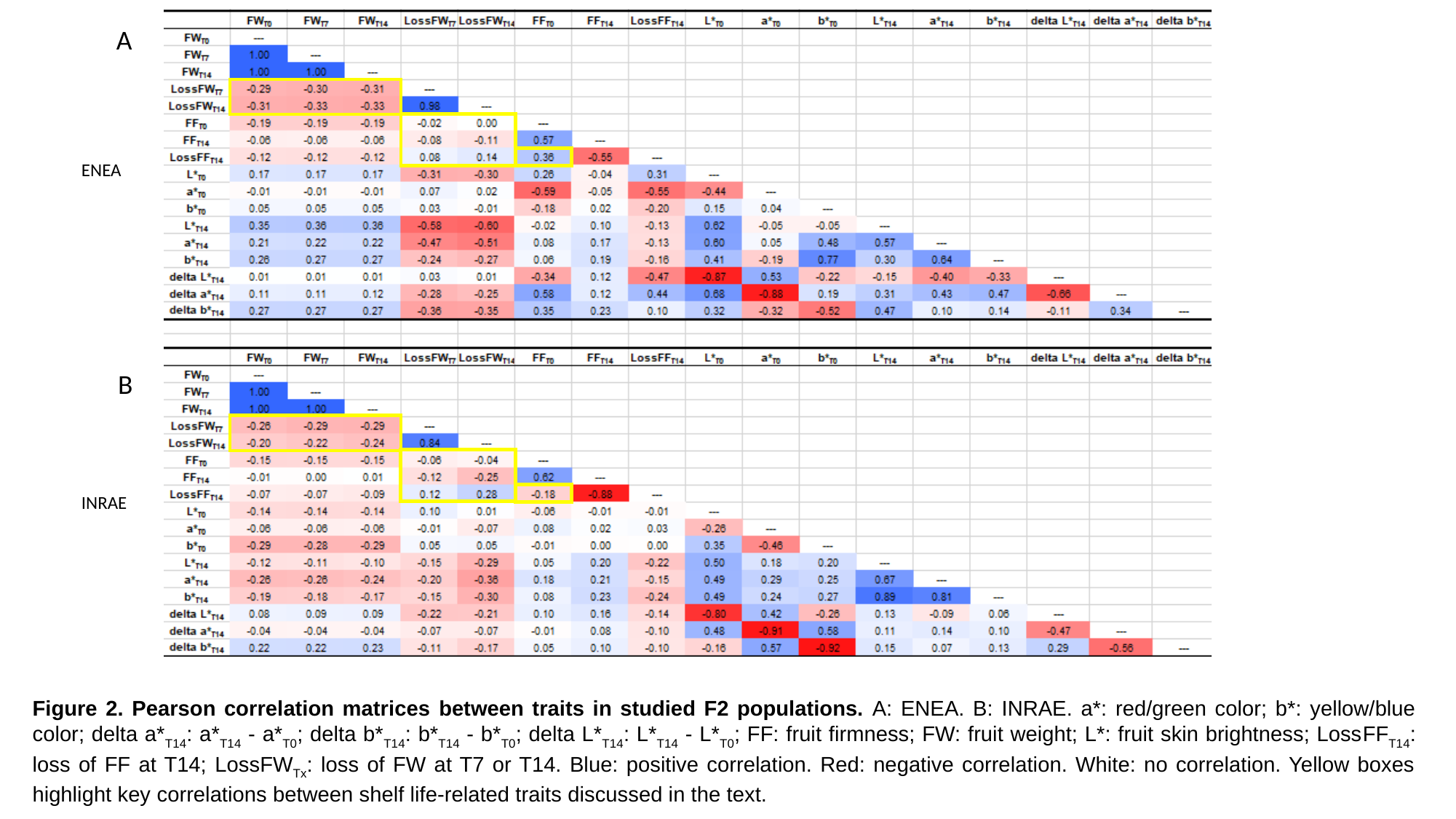

A
ENEA
B
INRAE
Figure 2. Pearson correlation matrices between traits in studied F2 populations. A: ENEA. B: INRAE. a*: red/green color; b*: yellow/blue color; delta a*_T14_: a*_T14_ - a*_T0_; delta b*_T14_: b*_T14_ - b*_T0_; delta L*_T14_: L*_T14_ - L*_T0_; FF: fruit firmness; FW: fruit weight; L*: fruit skin brightness; LossFF_T14_: loss of FF at T14; LossFW_Tx_: loss of FW at T7 or T14. Blue: positive correlation. Red: negative correlation. White: no correlation. Yellow boxes highlight key correlations between shelf life-related traits discussed in the text.

### Slide 4
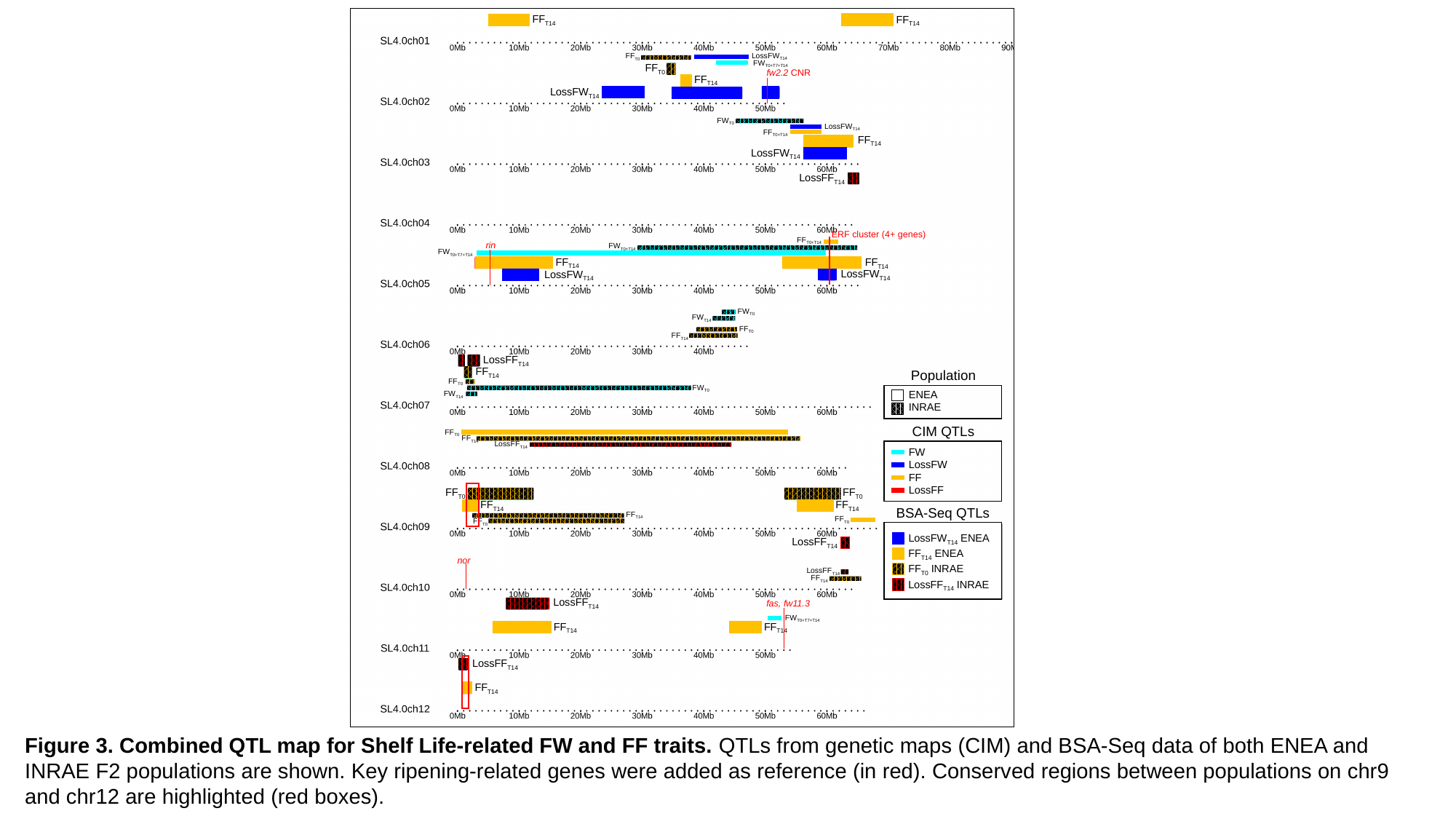

FF_T14_
FF_T14_
LossFW_T14_
FF_T0_
FW_T0+T7+T14_
FF_T0_
FF_T14_
LossFW_T14_
FW_T0_
LossFW_T14_
FF_T0+T14_
FF_T14_
LossFW_T14_
LossFF_T14_
FF_T0+T14_
FW_T0+T14_
FW_T0+T7+T14_
FF_T14_
FF_T14_
LossFW_T14_
LossFW_T14_
FW_T0_
FW_T14_
FF_T0_
FF_T14_
LossFF_T14_
FF_T14_
Population
FF_T0_
FW_T0_
ENEA
INRAE
FW_T14_
CIM QTLs
FF_T0_
FF_T14_
LossFF_T14_
FW
LossFW
FF
LossFF
FF_T0_
FF_T0_
FF_T14_
FF_T14_
BSA-Seq QTLs
FF_T14_
FF_T0_
FF_T0_
LossFW_T14_ ENEA
FF_T14_ ENEA
FF_T0_ INRAE
LossFF_T14_ INRAE
LossFF_T14_
nor
LossFF_T14_
FF_T14_
LossFF_T14_
FW_T0+T7+T14_
FF_T14_
FF_T14_
LossFF_T14_
FF_T14_
fw2.2 CNR
ERF cluster (4+ genes)
rin
fas, fw11.3
Figure 3. Combined QTL map for Shelf Life-related FW and FF traits. QTLs from genetic maps (CIM) and BSA-Seq data of both ENEA and INRAE F2 populations are shown. Key ripening-related genes were added as reference (in red). Conserved regions between populations on chr9 and chr12 are highlighted (red boxes).

### Slide 5
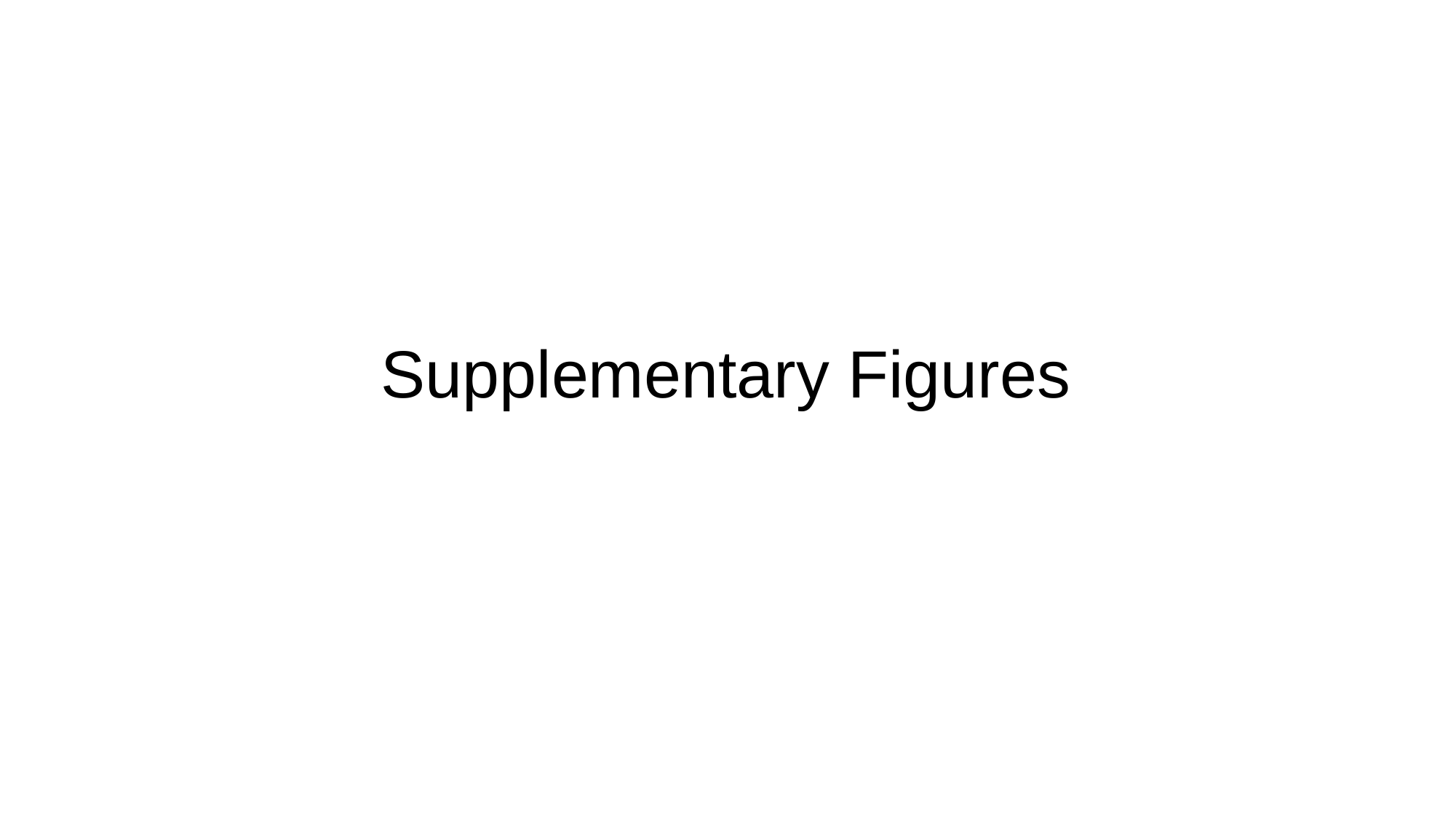

#### Supplementary Figures

### Slide 6
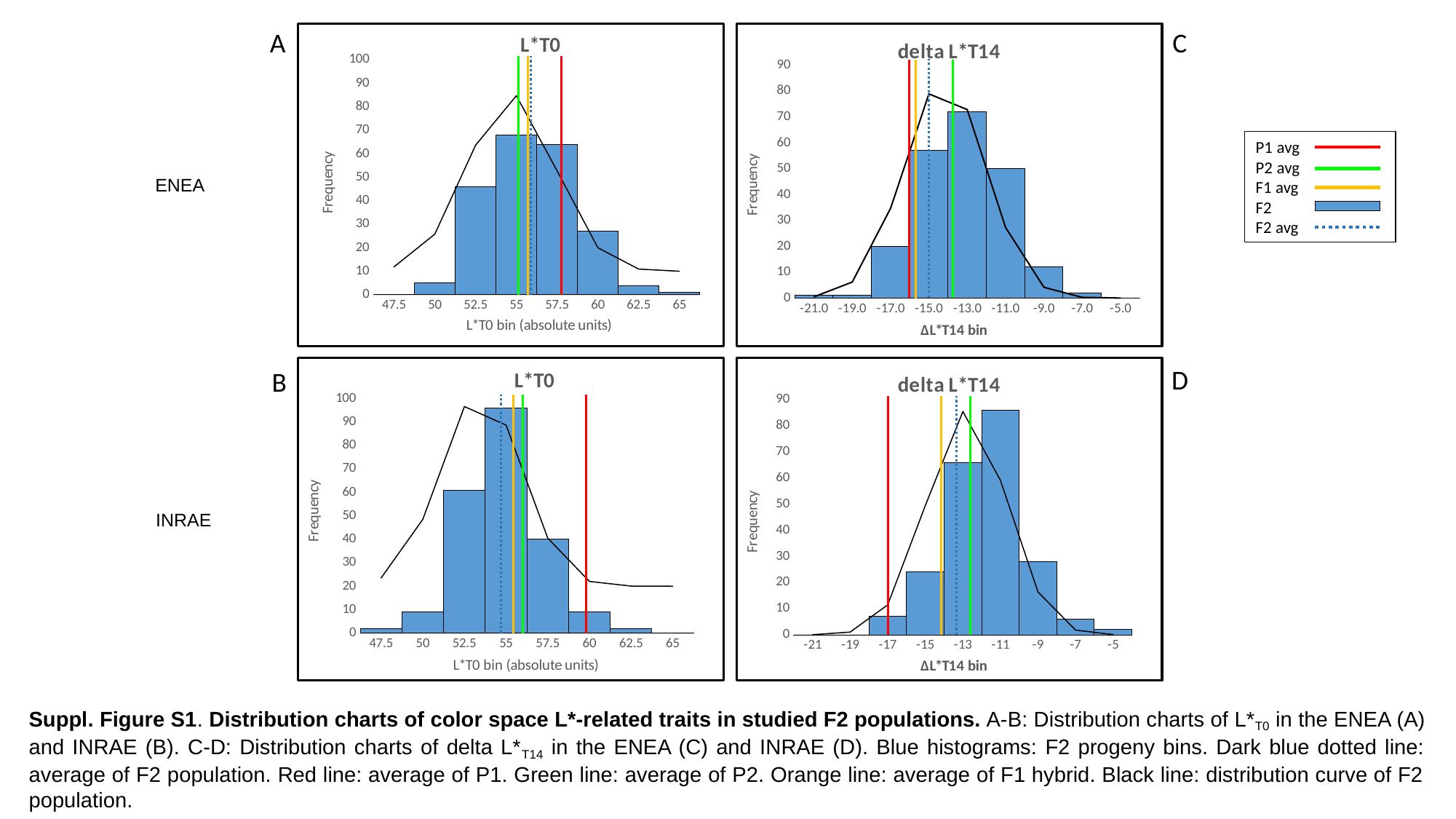

A
C
###### Chart: L*T0
| Category | | |
|---|---|---|
| 47.5 | 0.0 | 0.0038269016170385086 |
| 50 | 5.0 | 0.031636328113019196 |
| 52.5 | 46.0 | 0.10734381302536712 |
| 55 | 68.0 | 0.1494927524653821 |
| 57.5 | 64.0 | 0.08545066198899727 |
| 60 | 27.0 | 0.020047623526541693 |
| 62.5 | 4.0 | 0.0019304676927546607 |
| 65 | 1.0 | 7.629821028490264e-05 |
###### Chart: delta L*T14
| Category | | |
|---|---|---|
| -21 | 1.0 | 0.0009948788125155484 |
| -19 | 1.0 | 0.013753968893170565 |
| -17 | 20.0 | 0.07714233360980179 |
| -15 | 57.0 | 0.17553526772210243 |
| -13 | 72.0 | 0.1620477026374969 |
| -11 | 50.0 | 0.06069155287880718 |
| -9 | 12.0 | 0.00922190193860679 |
| -7 | 2.0 | 0.0005684858101795118 |
| -5 | 0.0 | 1.4217578563445165e-05 |
P1 avg
P2 avg
F1 avg
F2
F2 avg
ENEA
###### Chart: L*T0
| Category | | |
|---|---|---|
| 47.5 | 2.0 | 0.007060008719393219 |
| 50 | 9.0 | 0.05694472103230046 |
| 52.5 | 61.0 | 0.153134037039792 |
| 55 | 96.0 | 0.1372966763094738 |
| 57.5 | 40.0 | 0.04104103889969511 |
| 60 | 9.0 | 0.004090220279287676 |
| 62.5 | 2.0 | 0.00013590801807514616 |
| 65 | 0.0 | 1.5056134674827087e-06 |
###### Chart: delta L*T14
| Category | | |
|---|---|---|
| -21 | 0.0 | 8.739620943696582e-05 |
| -19 | 0.0 | 0.002358058567147857 |
| -17 | 7.0 | 0.025457473916647027 |
| -15 | 24.0 | 0.10997013462924937 |
| -13 | 66.0 | 0.19007847337999972 |
| -11 | 86.0 | 0.1314588543328717 |
| -9 | 28.0 | 0.03637855799706241 |
| -7 | 6.0 | 0.004028096316510404 |
| -5 | 2.0 | 0.00017846495306995686 |D
B
INRAE
Suppl. Figure S1. Distribution charts of color space L*-related traits in studied F2 populations. A-B: Distribution charts of L*_T0_ in the ENEA (A) and INRAE (B). C-D: Distribution charts of delta L*_T14_ in the ENEA (C) and INRAE (D). Blue histograms: F2 progeny bins. Dark blue dotted line: average of F2 population. Red line: average of P1. Green line: average of P2. Orange line: average of F1 hybrid. Black line: distribution curve of F2 population.

### Slide 7
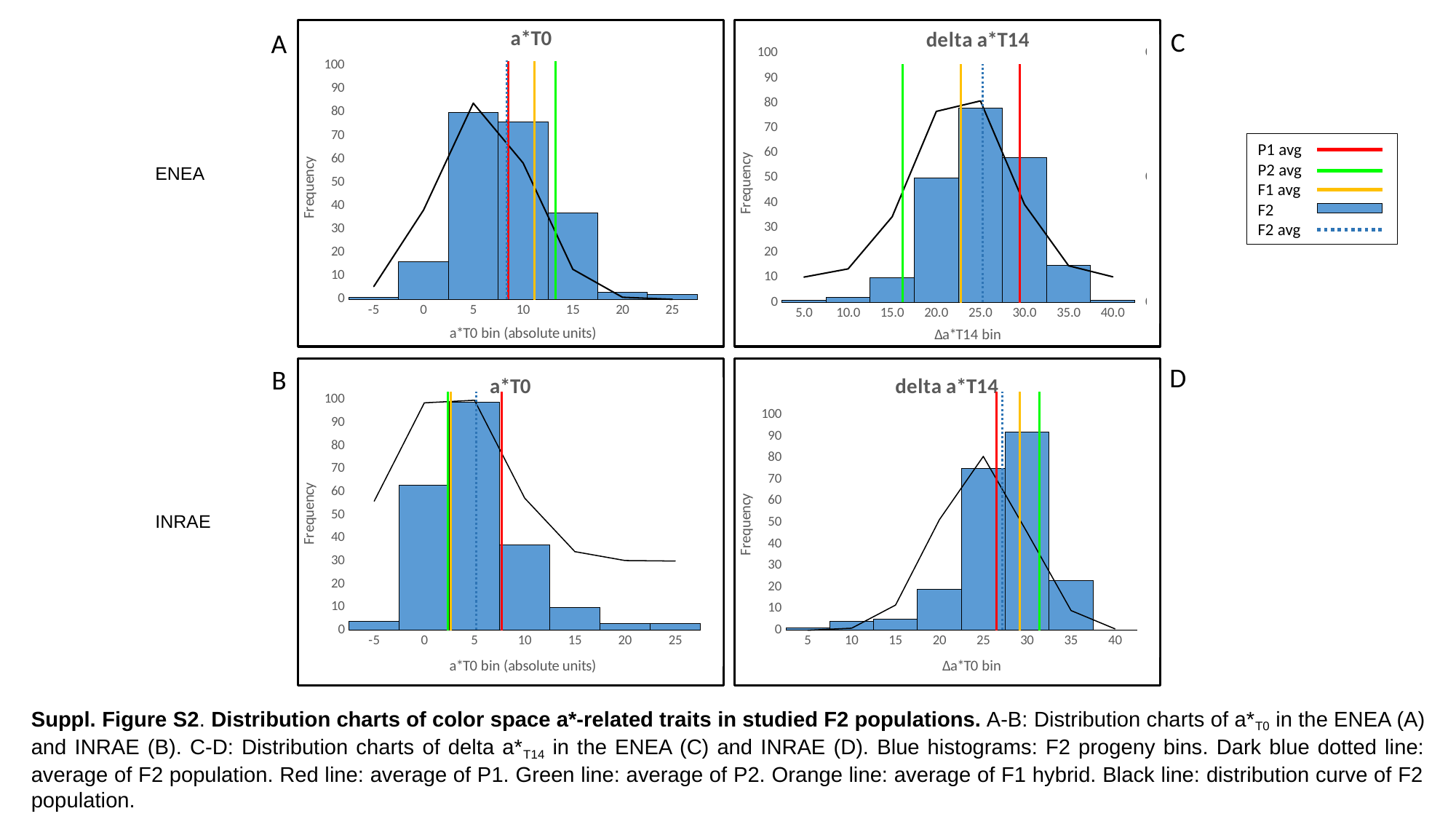

###### Chart: a*T0
| Category | | |
|---|---|---|
| -5 | 1.0 | 0.00551444599233772 |
| 0 | 16.0 | 0.03821422631972775 |
| 5 | 80.0 | 0.08389439284549388 |
| 10 | 76.0 | 0.05834793355139153 |
| 15 | 37.0 | 0.012855906476132162 |
| 20 | 3.0 | 0.0008973556660874114 |
| 25 | 2.0 | 1.9843176471978845e-05 |
###### Chart: delta a*T14
| Category | | |
|---|---|---|
| 5 | 1.0 | 0.00019260603803755843 |
| 10 | 2.0 | 0.0034755849357047948 |
| 15 | 10.0 | 0.024404403190533188 |
| 20 | 50.0 | 0.06667925726047393 |
| 25 | 78.0 | 0.07089173290821028 |
| 30 | 58.0 | 0.02932801761085572 |
| 35 | 15.0 | 0.004721196767471343 |
| 40 | 1.0 | 0.00029573571137897936 |C
A
P1 avg
P2 avg
F1 avg
F2
F2 avg
ENEA
D
B
###### Chart: a*T0
| Category | | |
|---|---|---|
| -5 | 4.0 | 0.025978696706739038 |
| 0 | 63.0 | 0.06863467401172337 |
| 5 | 99.0 | 0.06976129199101855 |
| 10 | 37.0 | 0.027279108893839096 |
| 15 | 10.0 | 0.00410384139652818 |
| 20 | 3.0 | 0.00023751753689339177 |
| 25 | 3.0 | 5.288658420735724e-06 |
###### Chart: delta a*T14
| Category | | |
|---|---|---|
| 5 | 1.0 | 2.7891402467374146e-05 |
| 10 | 4.0 | 0.0009546869015643894 |
| 15 | 5.0 | 0.011701293131755105 |
| 20 | 19.0 | 0.051355745466647965 |
| 25 | 75.0 | 0.08070984251052385 |
| 30 | 92.0 | 0.04541990751645458 |
| 35 | 23.0 | 0.009152679919958776 |
| 40 | 0.0 | 0.0006604388121846135 |
INRAE
Suppl. Figure S2. Distribution charts of color space a*-related traits in studied F2 populations. A-B: Distribution charts of a*_T0_ in the ENEA (A) and INRAE (B). C-D: Distribution charts of delta a*_T14_ in the ENEA (C) and INRAE (D). Blue histograms: F2 progeny bins. Dark blue dotted line: average of F2 population. Red line: average of P1. Green line: average of P2. Orange line: average of F1 hybrid. Black line: distribution curve of F2 population.

### Slide 8
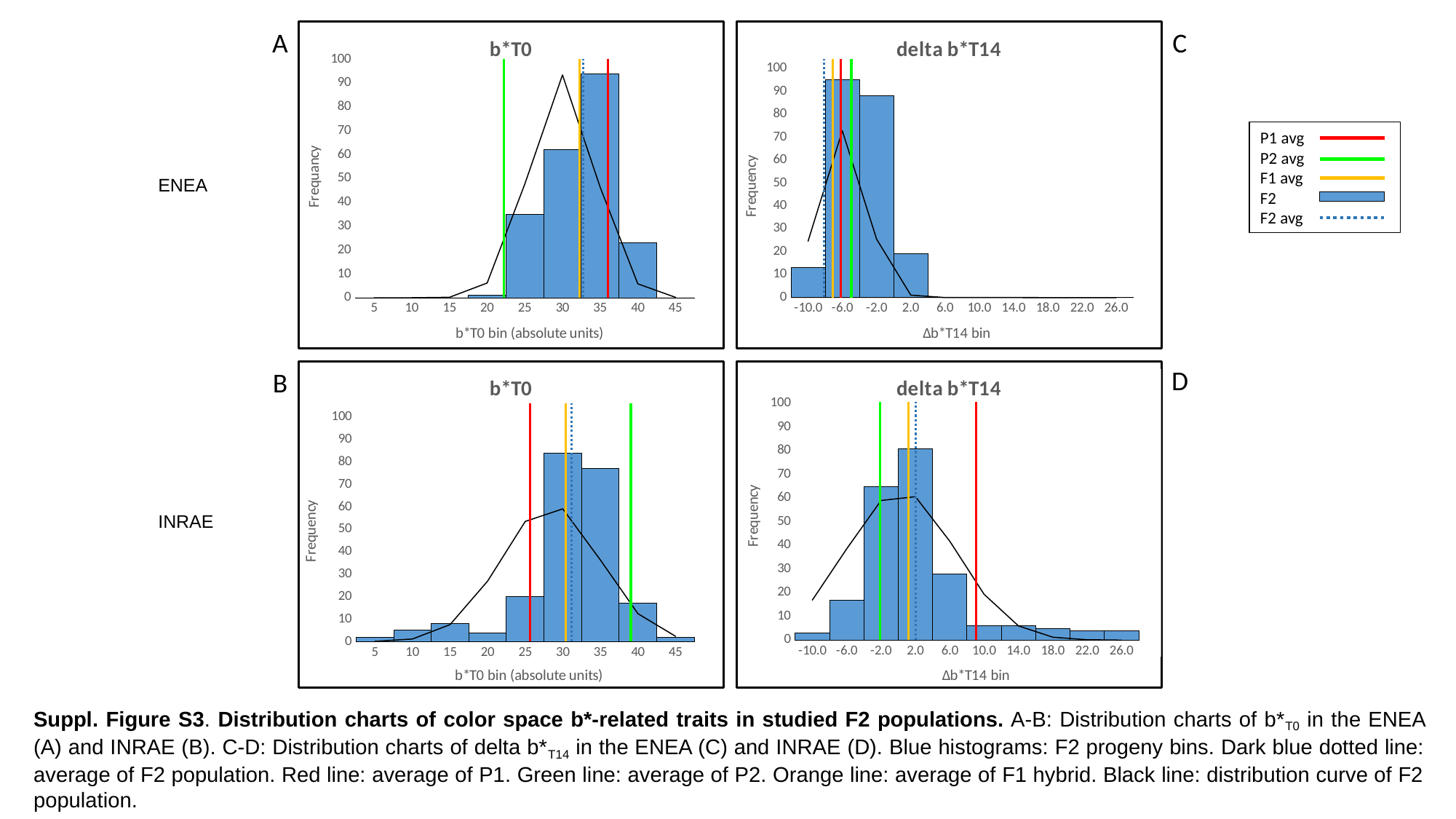

A
###### Chart: b*T0
| Category | | |
|---|---|---|
| 5 | 0.0 | 3.57375545545325e-09 |
| 10 | 0.0 | 1.6919272473370712e-06 |
| 15 | 0.0 | 0.00020312209797022923 |
| 20 | 1.0 | 0.0061837411026210605 |
| 25 | 35.0 | 0.04773798232426032 |
| 30 | 62.0 | 0.09345347523971169 |
| 35 | 94.0 | 0.046392255617326975 |
| 40 | 23.0 | 0.005840017616609437 |
| 45 | 0.0 | 0.00018642384521516575 |
###### Chart: delta b*T14
| Category | | |
|---|---|---|
| -10 | 13.0 | 0.04923810249999388 |
| -6 | 95.0 | 0.14565953405448234 |
| -2 | 88.0 | 0.051040412972487625 |
| 2 | 19.0 | 0.0021184933149475484 |
| 6 | 0.0 | 1.0415441191708834e-05 |
| 10 | 0.0 | 6.065490115540573e-09 |
| 14 | 0.0 | 4.184001027514201e-13 |
| 18 | 0.0 | 3.418655356702347e-18 |
| 22 | 0.0 | 3.308693313785563e-24 |
| 26 | 0.0 | 3.793109340900509e-31 |C
P1 avg
P2 avg
F1 avg
F2
F2 avg
ENEA
D
B
###### Chart: b*T0
| Category | | |
|---|---|---|
| 5 | 2.0 | 0.00010281763898429296 |
| 10 | 5.0 | 0.0011823943923056802 |
| 15 | 8.0 | 0.007569994525663963 |
| 20 | 4.0 | 0.026981571392313473 |
| 25 | 20.0 | 0.053539878648239767 |
| 30 | 84.0 | 0.05914608396303482 |
| 35 | 77.0 | 0.036375843946102396 |
| 40 | 17.0 | 0.012454853693883971 |
| 45 | 2.0 | 0.002374119920862892 |
###### Chart: delta b*T14
| Category | | |
|---|---|---|
| -10 | 3.0 | 0.016939627328947746 |
| -6 | 17.0 | 0.038592916479446175 |
| -2 | 65.0 | 0.05903747212399637 |
| 2 | 81.0 | 0.060640715017278864 |
| 6 | 28.0 | 0.041823201984394015 |
| 10 | 6.0 | 0.0193680836093104 |
| 14 | 6.0 | 0.006022439267357093 |
| 18 | 5.0 | 0.0012574033496485005 |
| 22 | 4.0 | 0.00017627600637833994 |
| 26 | 4.0 | 1.6593125525966183e-05 |
INRAE
Suppl. Figure S3. Distribution charts of color space b*-related traits in studied F2 populations. A-B: Distribution charts of b*_T0_ in the ENEA (A) and INRAE (B). C-D: Distribution charts of delta b*_T14_ in the ENEA (C) and INRAE (D). Blue histograms: F2 progeny bins. Dark blue dotted line: average of F2 population. Red line: average of P1. Green line: average of P2. Orange line: average of F1 hybrid. Black line: distribution curve of F2 population.

### Slide 9
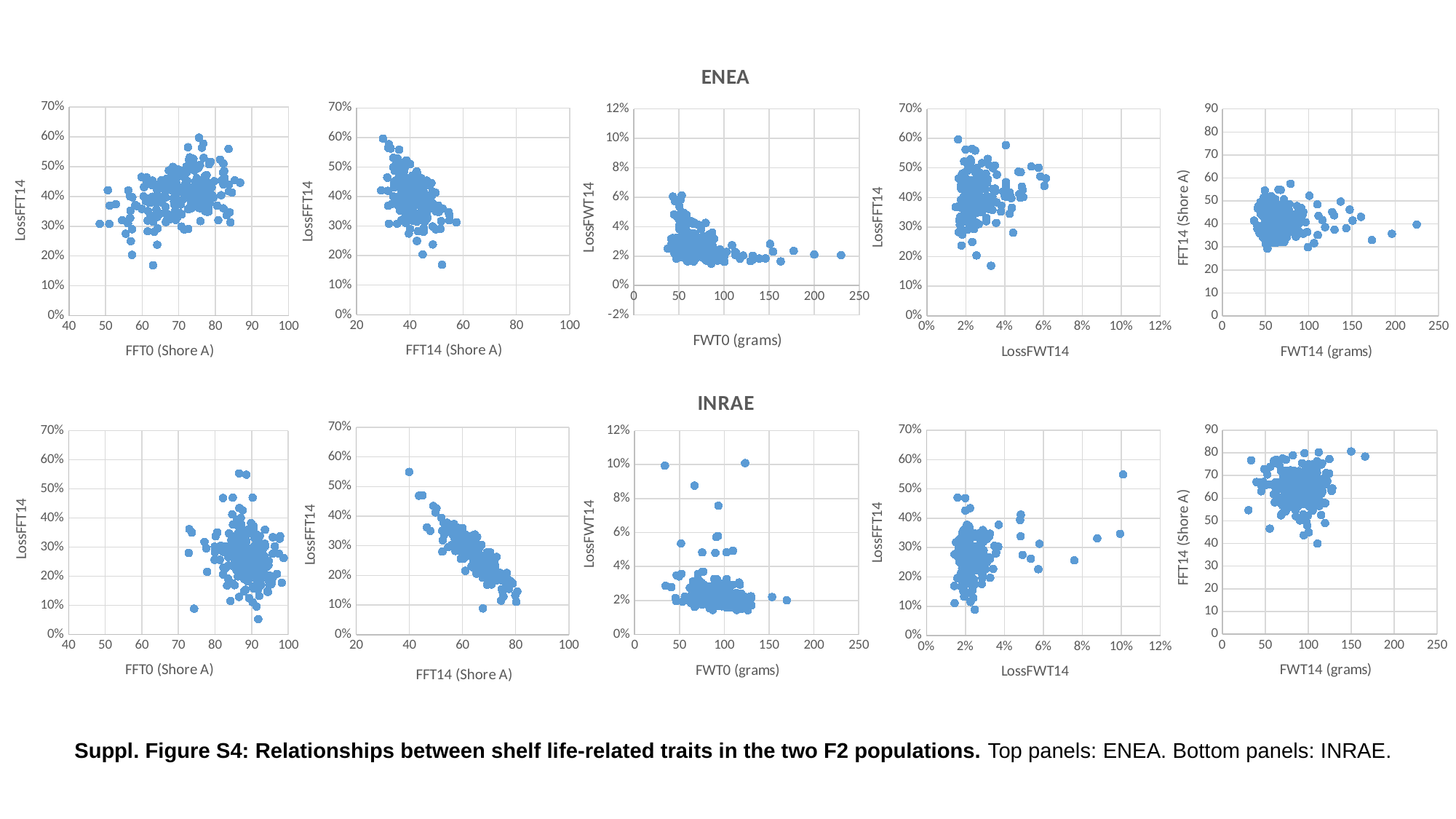

###### Chart: ENEA
| Category | LossFWT14 |
|---|---|
###### Chart
| Category | FFT14 |
|---|---|
###### Chart
| Category | ΔFFT14 |
|---|---|
###### Chart
| Category | LossFFT14 |
|---|---|
###### Chart
| Category | LossFFT14 |
|---|---|
###### Chart
| Category | ΔFFT14 |
|---|---|
###### Chart
| Category | FFT14 |
|---|---|
###### Chart: INRAE
| Category | LossFWT14 |
|---|---|
###### Chart
| Category | LossFFT14 |
|---|---|
###### Chart
| Category | LossFFT14 |
|---|---|Suppl. Figure S4: Relationships between shelf life-related traits in the two F2 populations. Top panels: ENEA. Bottom panels: INRAE.

### Slide 10
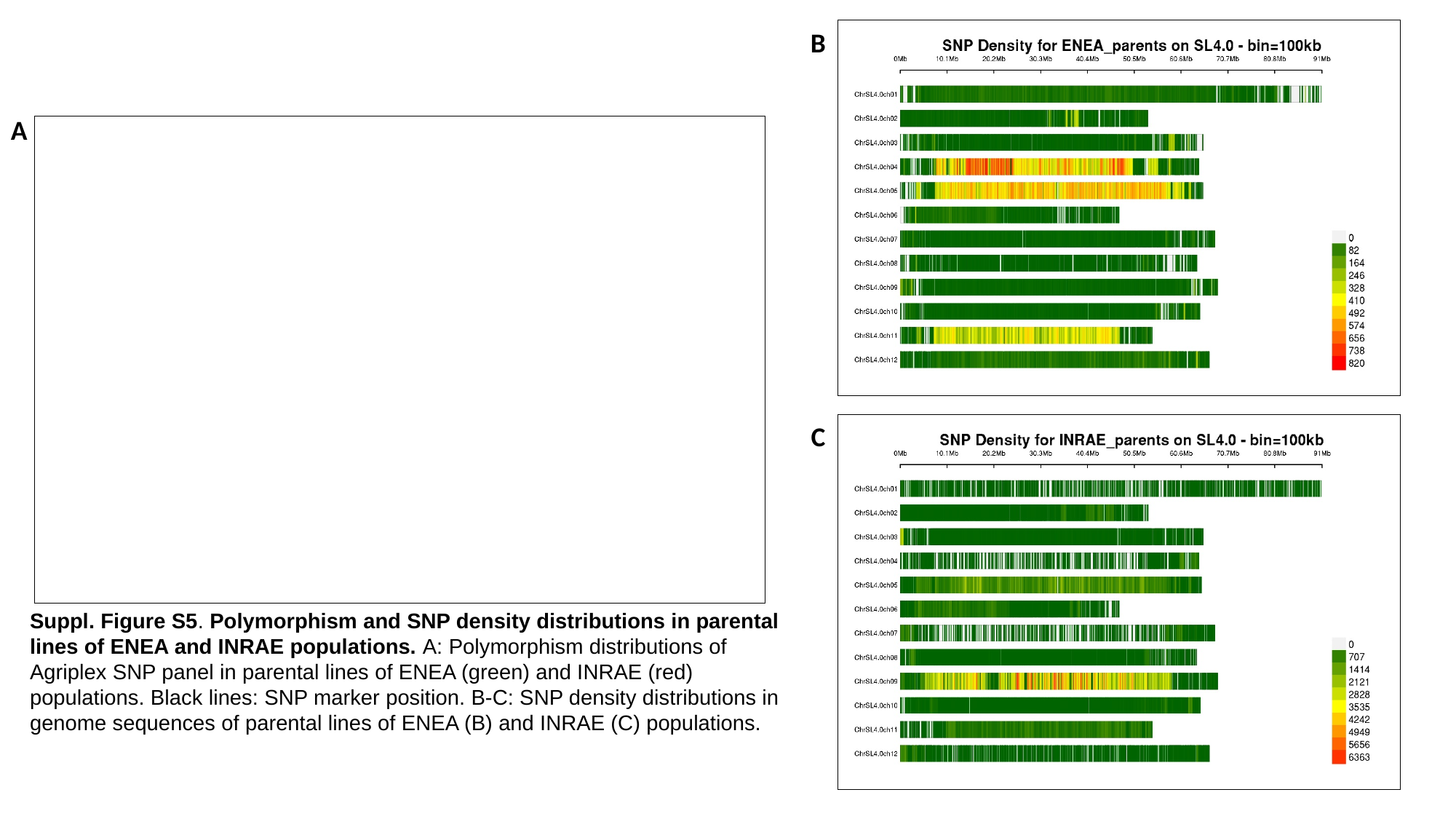

B
A
C
Suppl. Figure S5. Polymorphism and SNP density distributions in parental lines of ENEA and INRAE populations. A: Polymorphism distributions of Agriplex SNP panel in parental lines of ENEA (green) and INRAE (red) populations. Black lines: SNP marker position. B-C: SNP density distributions in genome sequences of parental lines of ENEA (B) and INRAE (C) populations.

### Slide 11
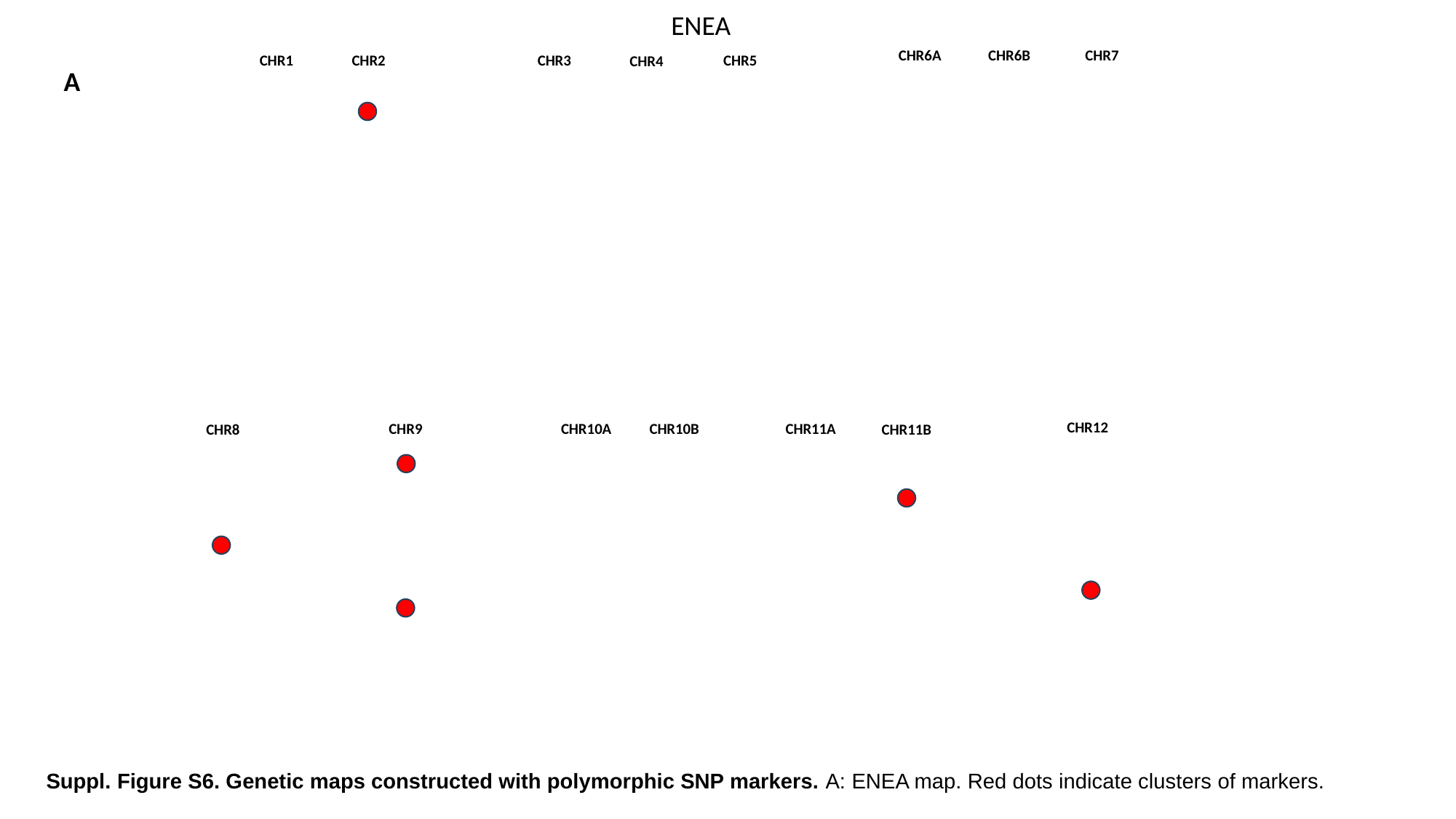

ENEA
CHR6A
CHR6B
CHR7
CHR1
CHR2
CHR3
CHR5
CHR4
A
CHR12
CHR11A
CHR10A
CHR10B
CHR9
CHR11B
CHR8
Suppl. Figure S6. Genetic maps constructed with polymorphic SNP markers. A: ENEA map. Red dots indicate clusters of markers.

### Slide 12
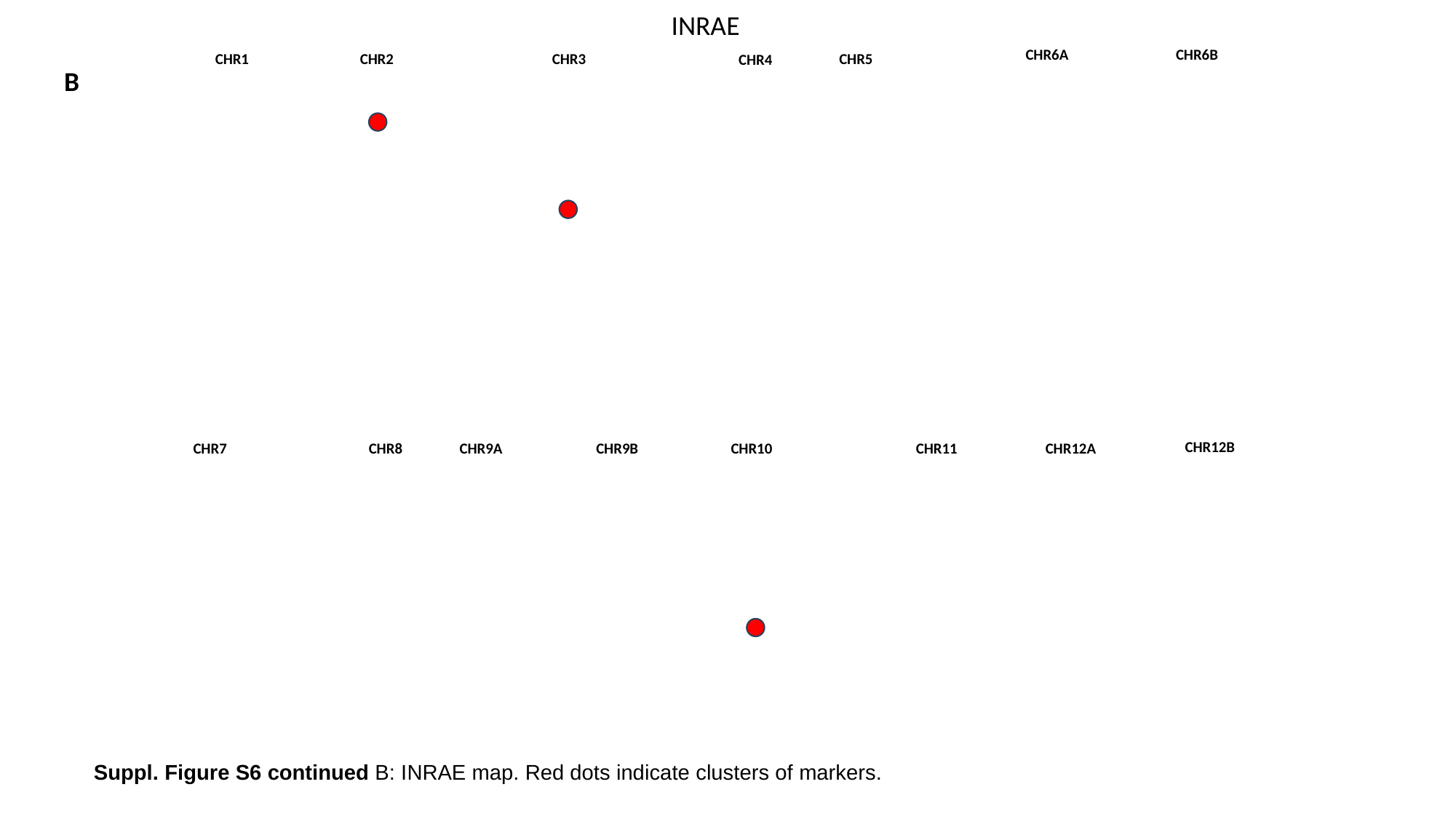

INRAE
CHR6A
CHR6B
CHR1
CHR2
CHR3
CHR5
CHR4
B
CHR12B
CHR11
CHR10
CHR9A
CHR12A
CHR7
CHR8
CHR9B
Suppl. Figure S6 continued B: INRAE map. Red dots indicate clusters of markers.

### Slide 13
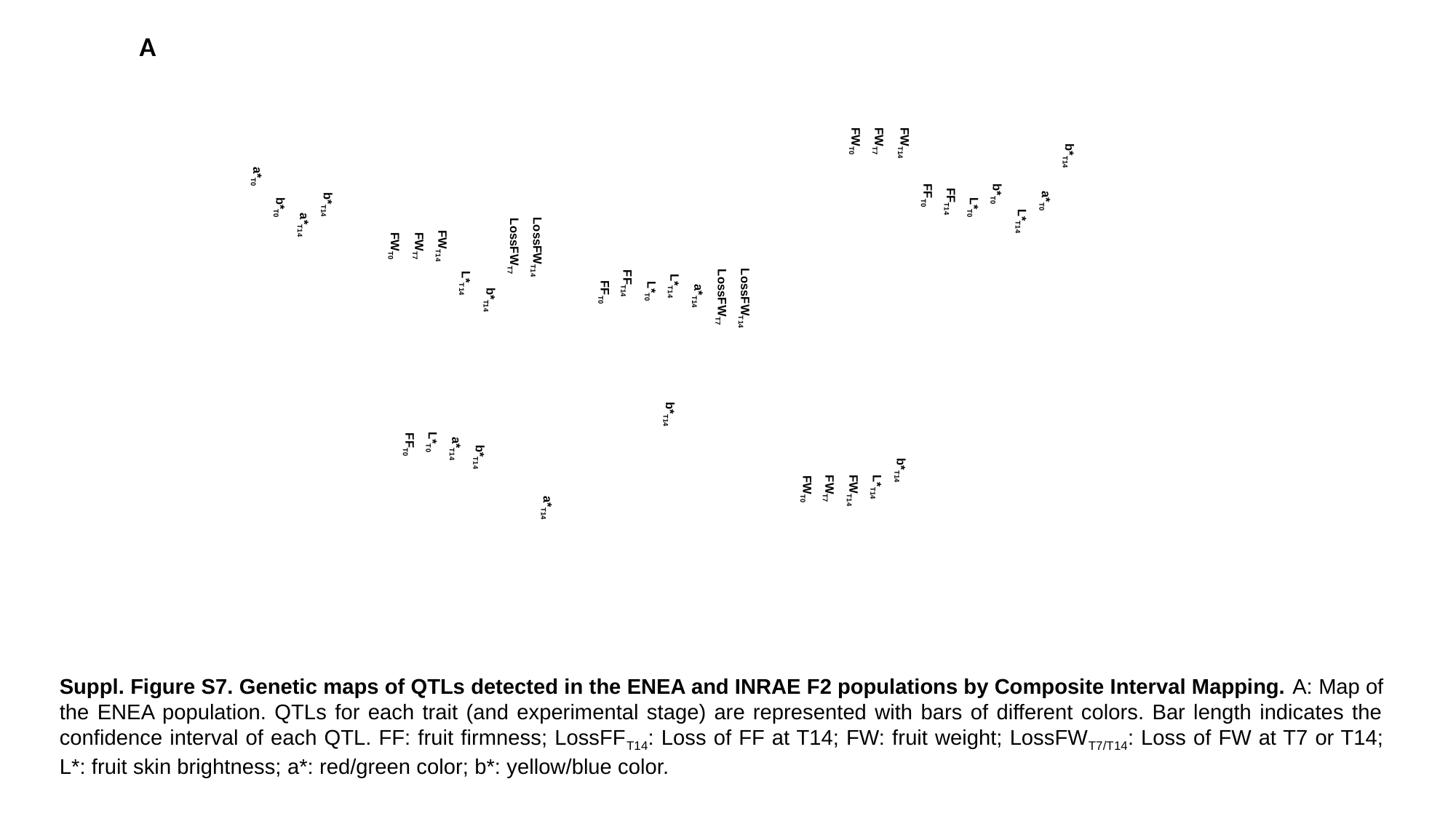

b*_T14_
a*_T0_
L*_T14_
b*_T0_
L*_T0_
FF_T14_
FF_T0_
FW_T14_
b*_T14_
FW_T7_
L*_T14_
FW_T0_
FW_T14_
FW_T7_
FW_T0_
LossFW_T14_
LossFW_T7_
a*_T14_
L*_T14_
b*_T14_
L*_T0_
FF_T14_
FF_T0_
a*_T14_
LossFW_T14_
LossFW_T7_
b*_T14_
b*_T14_
L*_T14_
a*_T14_
FW_T14_
L*_T0_
FW_T7_
FF_T0_
FW_T0_
b*_T14_
a*_T14_
b*_T0_
a*_T0_
A
Suppl. Figure S7. Genetic maps of QTLs detected in the ENEA and INRAE F2 populations by Composite Interval Mapping. A: Map of the ENEA population. QTLs for each trait (and experimental stage) are represented with bars of different colors. Bar length indicates the confidence interval of each QTL. FF: fruit firmness; LossFF_T14_: Loss of FF at T14; FW: fruit weight; LossFW_T7/T14_: Loss of FW at T7 or T14; L*: fruit skin brightness; a*: red/green color; b*: yellow/blue color.

### Slide 14
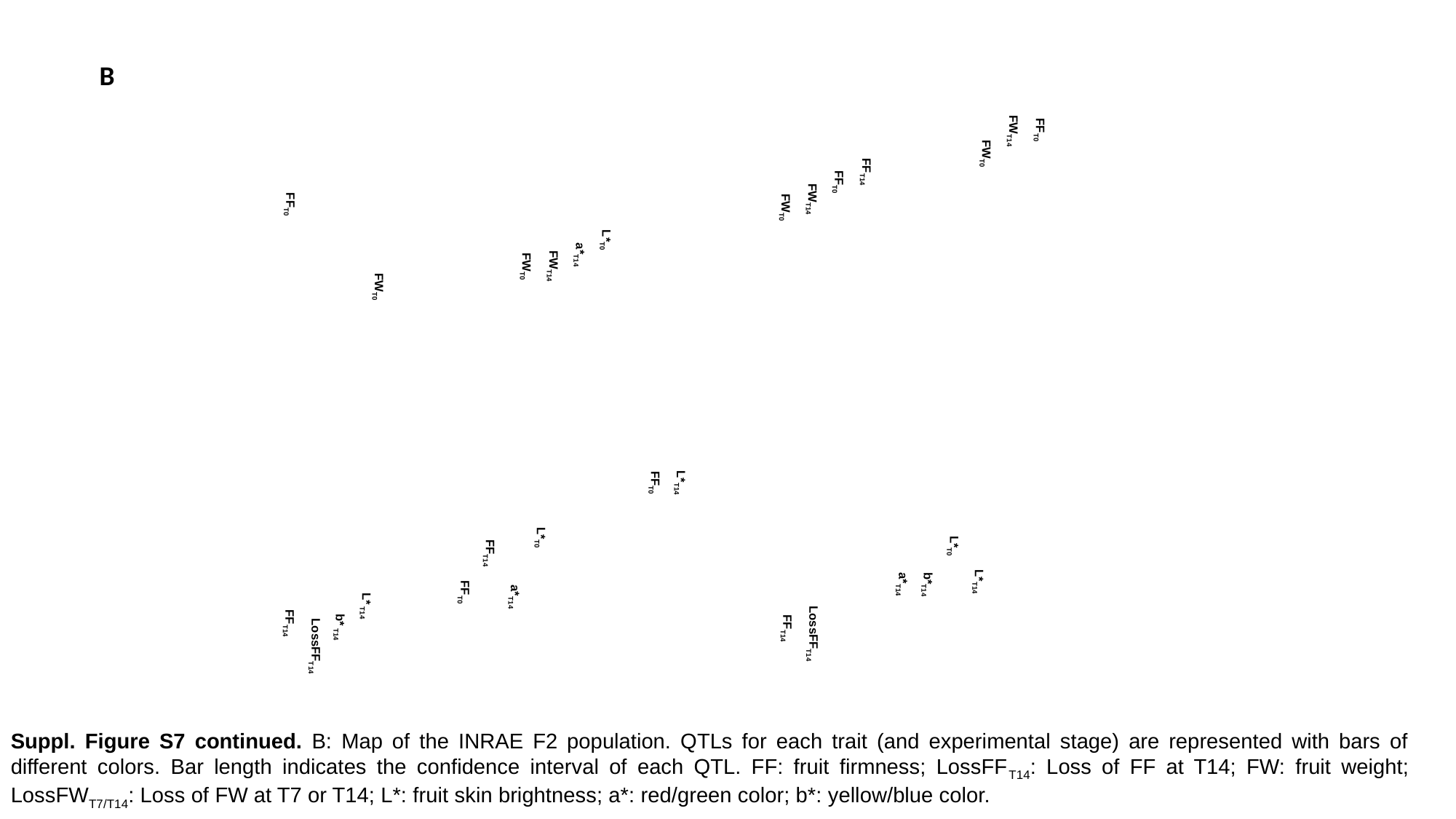

FF_T0_
FW_T14_
FW_T0_
L*_T14_
L*_T0_
b*_T14_
a*_T14_
FF_T14_
FF_T0_
LossFF_T14_
FW_T14_
FF_T14_
FW_T0_
L*_T14_
FF_T0_
L*_T0_
a*_T14_
FW_T14_
L*_T0_
FW_T0_
a*_T14_
FF_T14_
FF_T0_
FW_T0_
L* _T14_
b* _T14_
LossFF_T14_
FF_T0_
FF_T14_
B
Suppl. Figure S7 continued. B: Map of the INRAE F2 population. QTLs for each trait (and experimental stage) are represented with bars of different colors. Bar length indicates the confidence interval of each QTL. FF: fruit firmness; LossFF_T14_: Loss of FF at T14; FW: fruit weight; LossFW_T7/T14_: Loss of FW at T7 or T14; L*: fruit skin brightness; a*: red/green color; b*: yellow/blue color.

### Slide 15
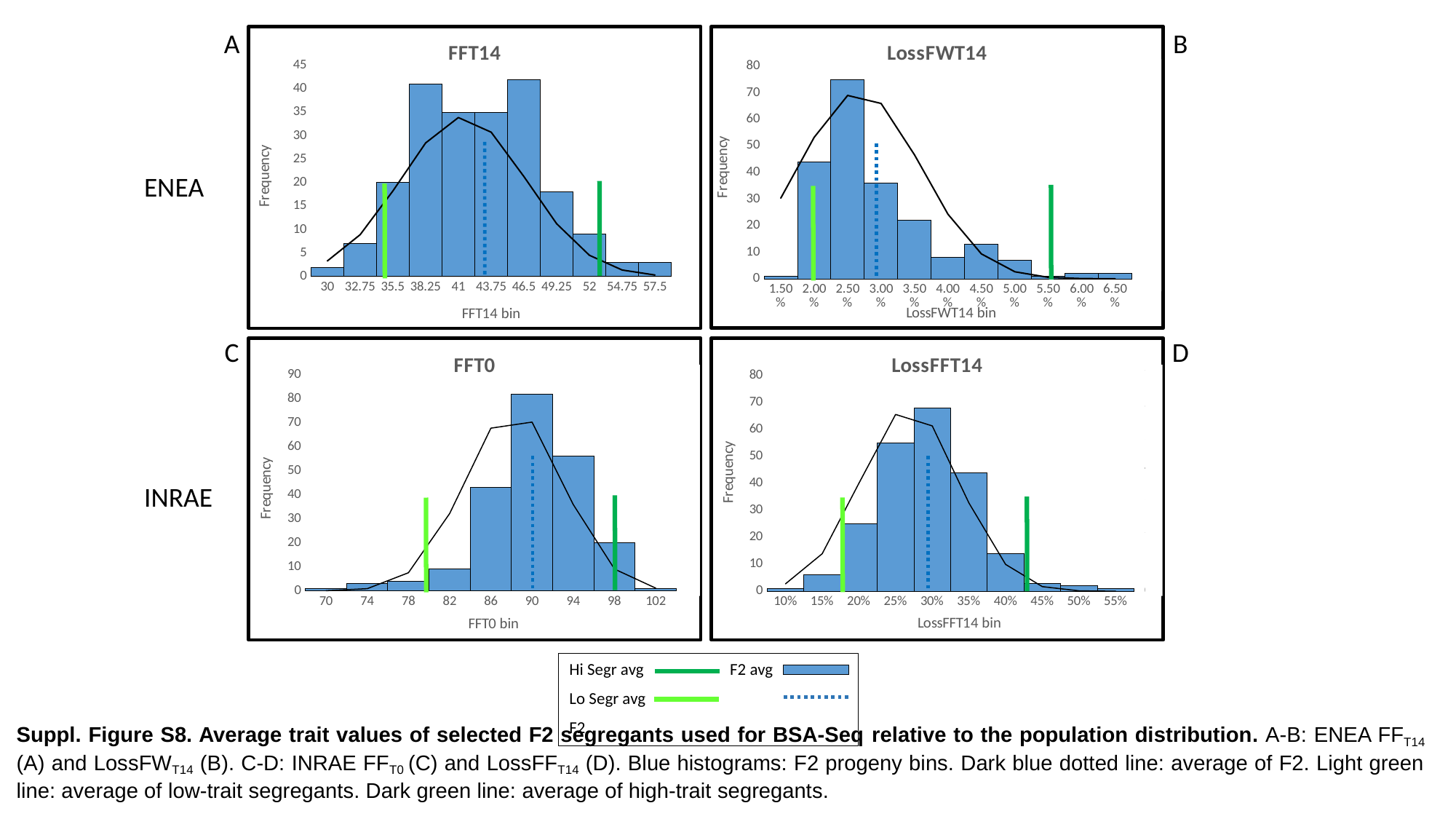

B
A
###### Chart: FFT14
| Category | | |
|---|---|---|
| 30 | 2.0 | 0.007387545862219576 |
| 32.75 | 7.0 | 0.01981519561203384 |
| 35.5 | 20.0 | 0.040538758697262874 |
| 38.25 | 41.0 | 0.0632581428038358 |
| 41 | 35.0 | 0.0752898334978158 |
| 43.75 | 35.0 | 0.06834868204779768 |
| 46.5 | 42.0 | 0.04732578876326401 |
| 49.25 | 18.0 | 0.02499421410659936 |
| 52 | 9.0 | 0.0100682738396182 |
| 54.75 | 3.0 | 0.003093459687655804 |
| 57.5 | 3.0 | 0.0007249495943393864 |
###### Chart: LossFWT14
| Category | | |
|---|---|---|
| 1.4999999999999999E-2 | 1.0 | 18.968642661259963 |
| 0.02 | 44.0 | 33.28012351355135 |
| 2.5000000000000001E-2 | 75.0 | 43.114394178343346 |
| 0.03 | 36.0 | 41.24281792380244 |
| 3.5000000000000003E-2 | 22.0 | 29.13151413936111 |
| 0.04 | 8.0 | 15.193802248662529 |
| 4.4999999999999998E-2 | 13.0 | 5.851383537665919 |
| 0.05 | 7.0 | 1.6639464509262702 |
| 5.5E-2 | 1.0 | 0.3493886641409363 |
| 0.06 | 2.0 | 0.05417101326642746 |
| 6.5000000000000002E-2 | 2.0 | 0.006201743388213101 |
ENEA
C
D
###### Chart: FFT0
| Category | | |
|---|---|---|
| 70 | 1.0 | 5.5087670187886e-05 |
| 74 | 3.0 | 0.0009679965637530389 |
| 78 | 4.0 | 0.008384171132373006 |
| 82 | 9.0 | 0.03579425775753089 |
| 86 | 43.0 | 0.07532402108996836 |
| 90 | 82.0 | 0.07813048849851847 |
| 94 | 56.0 | 0.039946109789098454 |
| 98 | 20.0 | 0.010066890649691781 |
| 102 | 1.0 | 0.001250498339220805 |
###### Chart: LossFFT14
| Category | | |
|---|---|---|
| 0.1 | 1.0 | 0.24231037683962833 |
| 0.15 | 6.0 | 1.2192005043923428 |
| 0.2 | 25.0 | 3.500996324309289 |
| 0.25 | 55.0 | 5.737485175461461 |
| 0.3 | 68.0 | 5.3661758262444454 |
| 0.35 | 44.0 | 2.8643205648172323 |
| 0.4 | 14.0 | 0.8725528555489311 |
| 0.45 | 3.0 | 0.15169639765343668 |
| 0.5 | 2.0 | 0.015051236910152631 |
| 0.55000000000000004 | 1.0 | 0.0008522804297717059 |
INRAE
Hi Segr avg
Lo Segr avg
F2
F2 avg
Suppl. Figure S8. Average trait values of selected F2 segregants used for BSA-Seq relative to the population distribution. A-B: ENEA FF_T14_ (A) and LossFW_T14_ (B). C-D: INRAE FF_T0_ (C) and LossFF_T14_ (D). Blue histograms: F2 progeny bins. Dark blue dotted line: average of F2. Light green line: average of low-trait segregants. Dark green line: average of high-trait segregants.

### Slide 16
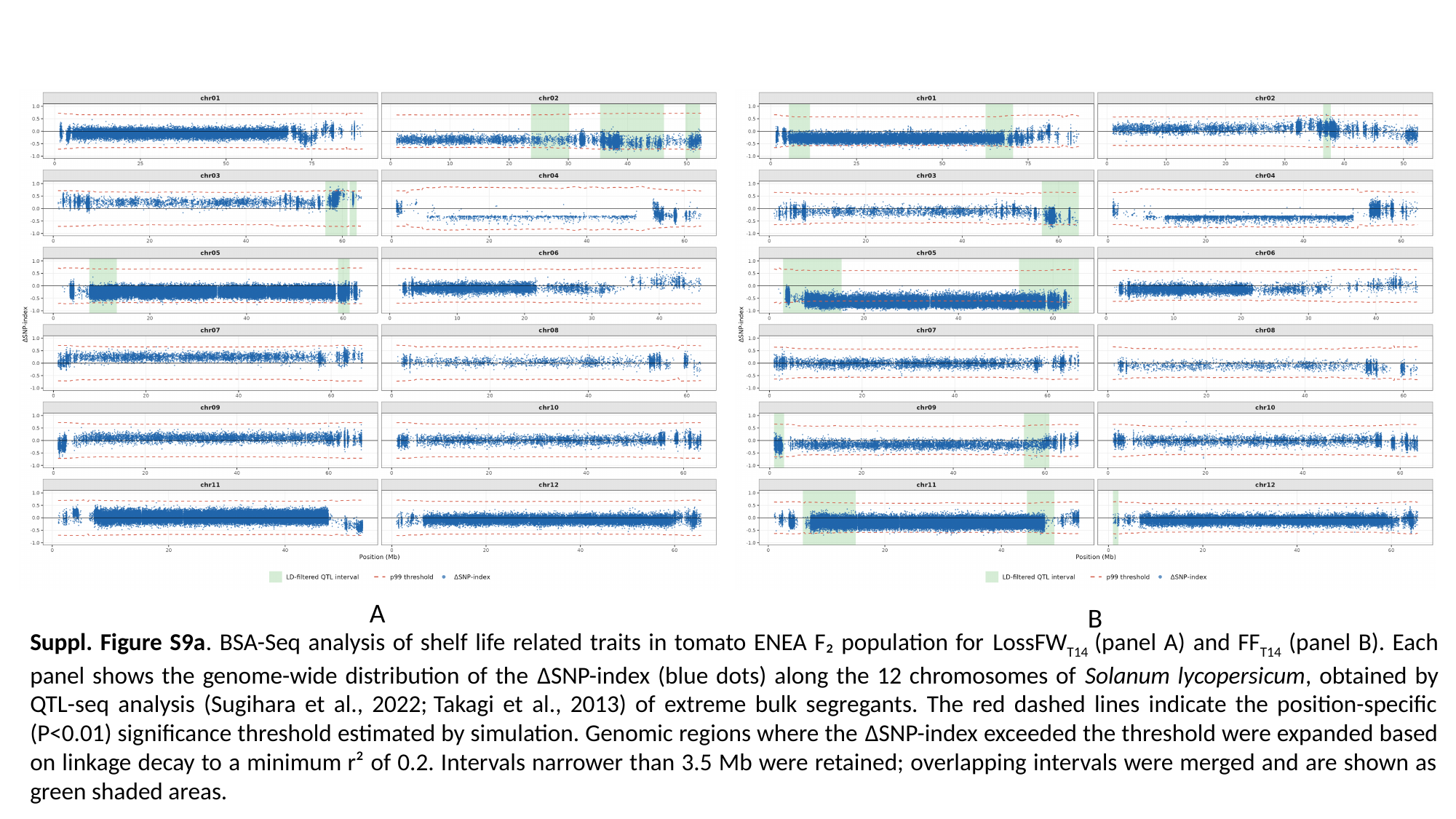

A
B
Suppl. Figure S9a. BSA-Seq analysis of shelf life related traits in tomato ENEA F₂ population for LossFW_T14_ (panel A) and FF_T14_ (panel B). Each panel shows the genome-wide distribution of the ΔSNP-index (blue dots) along the 12 chromosomes of Solanum lycopersicum, obtained by QTL-seq analysis (Sugihara et al., 2022; Takagi et al., 2013) of extreme bulk segregants. The red dashed lines indicate the position-specific (P<0.01) significance threshold estimated by simulation. Genomic regions where the ΔSNP-index exceeded the threshold were expanded based on linkage decay to a minimum r² of 0.2. Intervals narrower than 3.5 Mb were retained; overlapping intervals were merged and are shown as green shaded areas.

### Slide 17
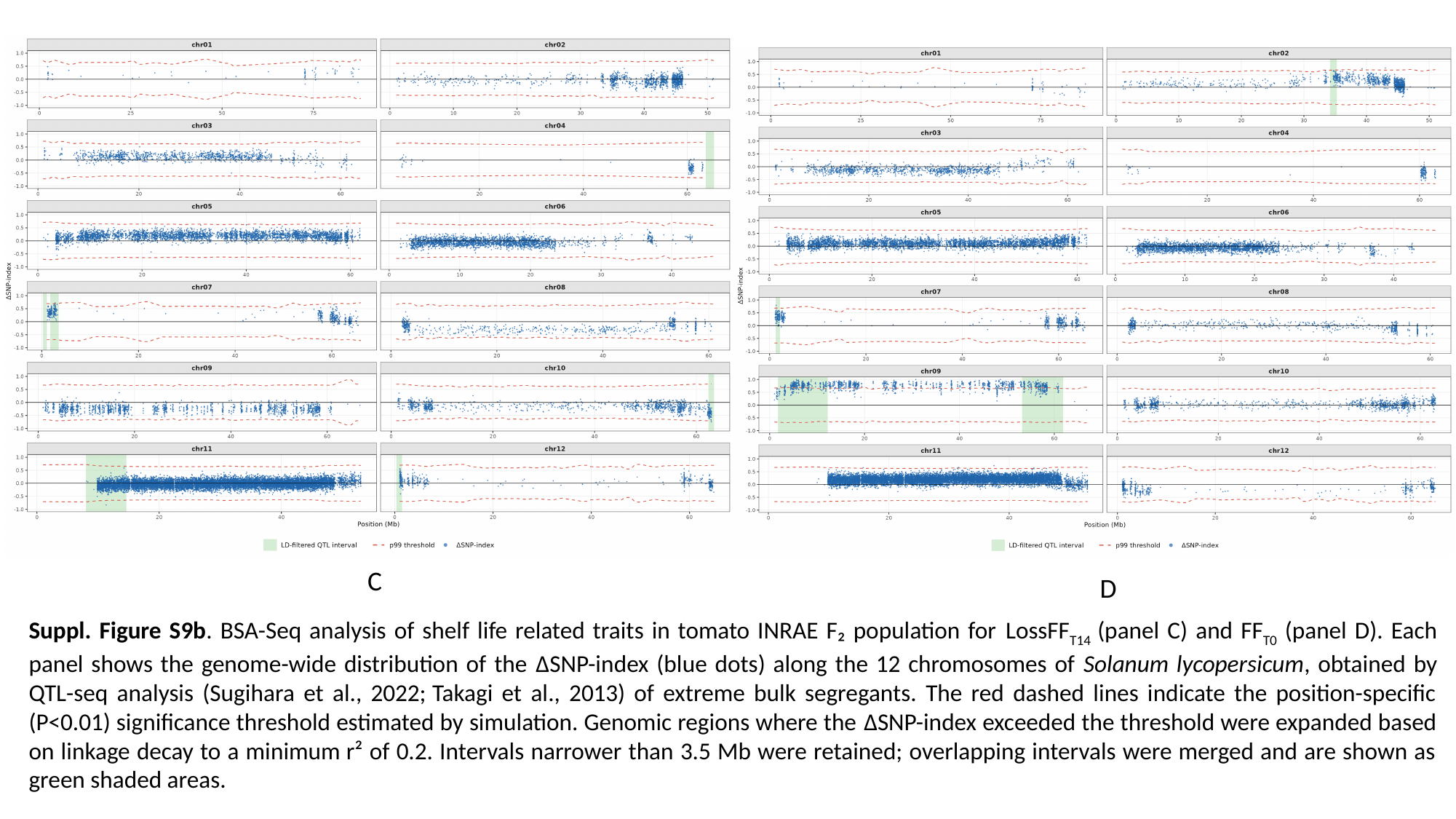

C
D
Suppl. Figure S9b. BSA-Seq analysis of shelf life related traits in tomato INRAE F₂ population for LossFF_T14_ (panel C) and FF_T0_ (panel D). Each panel shows the genome-wide distribution of the ΔSNP-index (blue dots) along the 12 chromosomes of Solanum lycopersicum, obtained by QTL-seq analysis (Sugihara et al., 2022; Takagi et al., 2013) of extreme bulk segregants. The red dashed lines indicate the position-specific (P<0.01) significance threshold estimated by simulation. Genomic regions where the ΔSNP-index exceeded the threshold were expanded based on linkage decay to a minimum r² of 0.2. Intervals narrower than 3.5 Mb were retained; overlapping intervals were merged and are shown as green shaded areas.
